# A chorionic gonadotropin assay enables non-invasive detection of ovulation and early pregnancy in a New World primate model

**DOI:** 10.64898/2026.03.12.711492

**Authors:** Keiko Kishimoto, Takuma Soga, Akio Iio, Masahiko Hatakeyama, Satoru Kawai, Michiko Kamioka, Jinsho Aoki, Yuka Bunzui, Yuko Yamada, Miho Kohara, Yoko Kurotaki, Wakako Kumita, Julie Brent-Cummins, Sang Su Oh, Milton Herrera, Lian Bik, Heather Narver, Tadashi Sankai, Tomoji Mashimo, Kazumasa Fukasawa, Erika Sasaki

**Author notes:** **Corresponding author:** Erika Sasaki, Division of Advanced Physiology, Central Institute for Experimental Medicine and Life Science (CIEM), 3-25-12 Tonomachi, Kawasaki-ku, Kawasaki, Kanagawa, 216-0821, Japan. These authors contributed equally to this work (Keiko Kishimoto, Takuma Soga).

## Abstract

Early detection of ovulation and pregnancy in the common marmoset is crucial for reproductive studies, yet hCG kits lack cross-reactivity with marmoset CG, and current methods remain labor-intensive. Here, we developed monoclonal antibodies against marmoset CGα and CGβ, and established a non-invasive immunochromatographic CG assay. By eliminating invasive blood sampling, this assay supports 3Rs principles and enables practical endocrine monitoring. The assay detected urinary CG surges preceding ovulation, enabling efficient embryo recovery through artificial insemination (75%). Early pregnancy was detected at approximately 17 days post-ovulation. In addition, pregnancy detection in squirrel monkeys suggests conservation of CG features among certain New World primates. Overall, this simple, non-invasive assay provides a practical tool for marmoset research and establishes a foundation for future conservation-oriented reproductive monitoring following appropriate species-specific validation.

## Introduction

The common marmoset (marmoset, *Callithrix jacchus*) is a New World monkey, native to the northeastern coast of Brazil, and has become an important model in biomedical research owing to its physiological similarities to humans, small body size, and the availability of genetic modification technologies. Within a research environment increasingly shaped by New Approach Methodologies (NAMs), marmosets remain indispensable in contexts where organism-level validation is required, including reproductive, developmental, and genetic modification studies. In these settings, reliable early detection of ovulation and pregnancy is essential for reproductive research, embryogenesis studies, the generation of genetically modified marmosets, and the optimization of breeding programs.

Chorionic gonadotropin (CG) is a crucial hormone involved in the establishment and maintenance of pregnancy, primarily through its roles in placentation and early embryonic development ^1^. CG belongs to the glycoprotein hormone family, which also includes luteinizing hormone (LH), thyroid-stimulating hormone (TSH), and follicle-stimulating hormone (FSH). Members of this family are heterodimeric glycoproteins composed of a common α subunit and hormone-specific β subunits that confer distinct biological functions ^2,3^.

LH and CG act through the same luteinizing hormone receptor (LHR) ^4,5^, which consists of 11 exons and 10 introns at the genomic level ^6^. Müller et al. demonstrated that deletion of exon 10 in the LHR (LHR-ex10) severely impairs LH-mediated signal transduction, whereas CG-mediated signaling remains functional ^7,8^. Notably, New World monkeys, including marmosets, lack exon 10 in the mature LHR protein, despite its presence at the genomic level ^9,10^. Furthermore, Gromoll et al. reported that CG is highly expressed in the marmoset pituitary and that the absence of exon 10 may compromise normal LH bioactivity across New World primates ^10^. Supporting this, Müller et al. reported that only CGβ subunit mRNA, but not LHβ, is expressed in the marmoset pituitary, suggesting that a CG surge, rather than an LH surge, drives late follicular maturation and ovulation in New World monkeys.

In humans, urinary CG immunochromatographic test kits based on antibodies against the CGβ subunit are widely used for pregnancy detection. Although the human and marmoset CGβ subunits share high overall sequence similarity, their amino acid identity is substantially lower, resulting in a lack of cross-reactivity between human CG immunochromatographic test kits and marmoset urine. Consequently, pregnancy in marmosets is currently confirmed by sustained elevation of plasma progesterone levels over several weeks and through uterine ultrasonographic examination. These approaches require repeated animal handling for blood collection or imaging, are labor-intensive, and impose stress on the animals, making them undesirable from an animal welfare perspective.

In this study, we generated monoclonal antibodies targeting marmoset CGα and CGβ and developed a marmoset-validated immunochromatographic CG assay (CG assay) for ovulation monitoring and early pregnancy detection. Using this marmoset CG assay, we aimed to enable precise estimation of ovulation, efficient embryo recovery via artificial insemination, and early pregnancy detection. This advancement is particularly significant given the emerging evidence of fundamental differences in early embryonic development between rodents and primates, including humans. By precisely estimating ovulation and achieving high embryo recovery rates, our CG assay provides a crucial foundation for accelerating research on early primate embryonic development.

## Results

### Detection of ovulation and CG surge

We confirmed the detection limit of the developed CG assay. As shown in Fig. 1 and Table 1; the test line reliably detected CG concentrations 6.3 AU/mL or above. A CG score of “3” was assigned to urine CG concentration between 6.3 AU/mL and 12.5 AU/mL, with scores 3 or above defined as positive. All subsequent experiments were performed according to this definition.

**Fig. 1.**
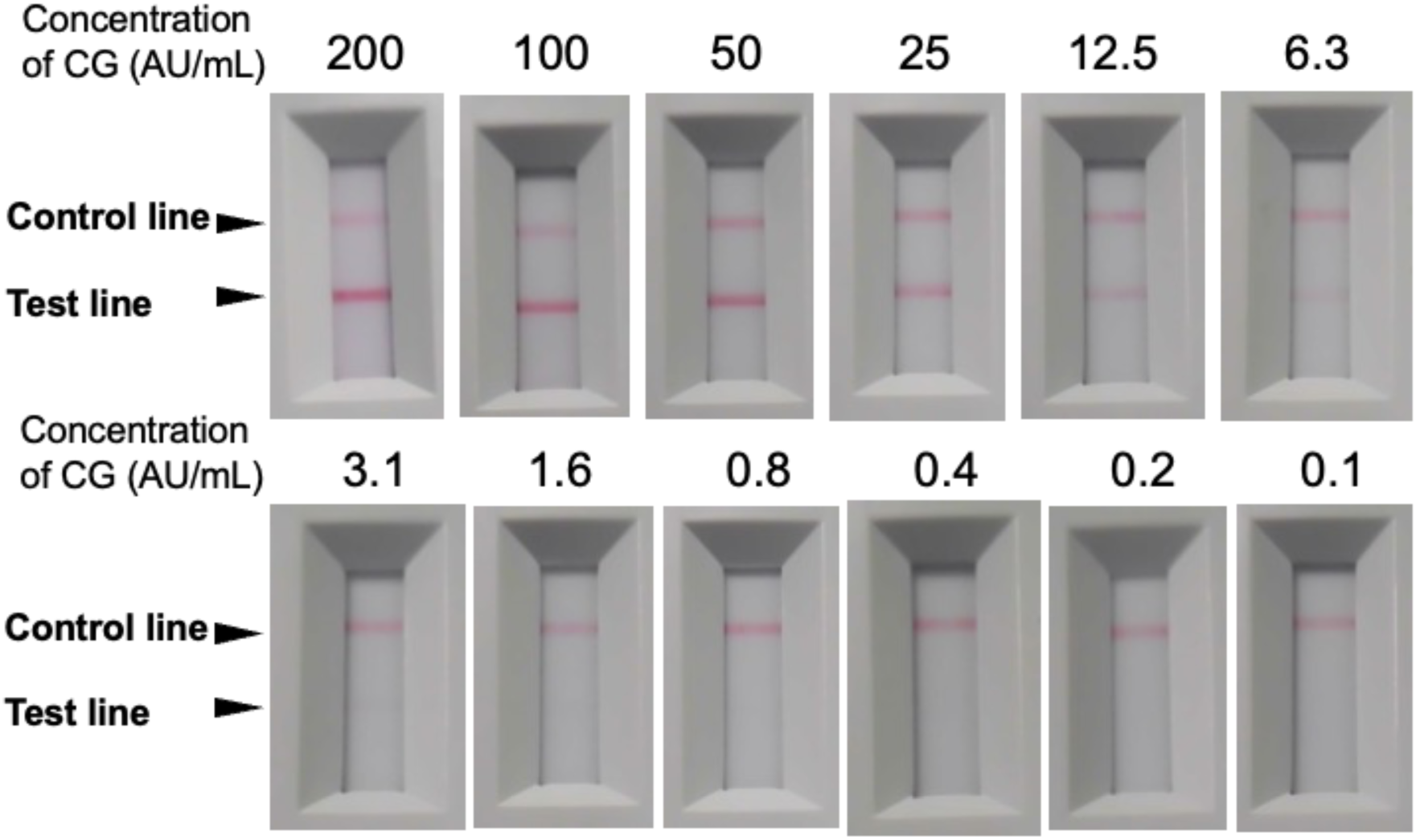
The relationship between test line density and CG concentration. The relationship between test line density and CG concentration was decided. The upper line is the control line, and the lower line is the test line. CG concentrations are summarized on Table 1. The higher the concentration, the clearer the test line was detected.

**Table 1.** The relationship of CG concentration and score.

| Concentration of CG (AU/mL) | 200 | 100 | 50 | 25 | 12.5 | 6.3 | 3.1 | 1.6 | 0.8 | 0.4 | 0.2 | 0.1 |
| --- | --- | --- | --- | --- | --- | --- | --- | --- | --- | --- | --- | --- |
| Results | + | + | + | + | + | + | ± | ± | ± | — | — | — |
| Score | 5 | 5 | 4 | 4 | 3 | 3 | 2 | 2 | 2 | 1 | 1 | 1 |

To verify the effectiveness of the CG assay, the endocrine profiles that regulate the ovarian cycle were assessed. Plasma concentrations of progesterone (P4) and estradiol (E2) were measured using enzyme immunoassay (EIA), while urinary concentrations of CG were measured using Enzyme-Linked Immunosorbent Assay (ELISA). The ELISA-derived urinary CG concentrations were compared with the CG scores.

In all animals, an E2 surge (plasma E2 concentration: ≥435.6 pg/mL; mean: 699 ± 274.69 pg/ml) was observed at 6.5 ± 1.0 days after prostaglandin F2α (PGF2α) administration. Plasma progesterone levels subsequently rose above 10.0 ng/mL at 9.25 ± 1.5 days. In addition, the urinary CG concentration was increased to an average of 224.29 ± 134.82 AU/mL at 8.0 ± 1.41 days after the PGF2α injection, and at 1.50 ± 1.0 days after the E2 surge (Fig. 2). Further, the CG scores exceeded 3 at an average of 7.5± 1.0 days after the PGF2α injection (1.33 ± 1.15 days after the E2 surge), peaking at 8.0 ± 1.41 days after the PGF2α injection (1.5 ± 1.29 days after the E2 surge). These findings indicate that after PGF2α administration, plasma E2 concentration temporarily increased prior to urine CG level, followed by plasma P4 levels. These endocrine profiles were consistent with other New World species ^11^.

**Fig. 2.**
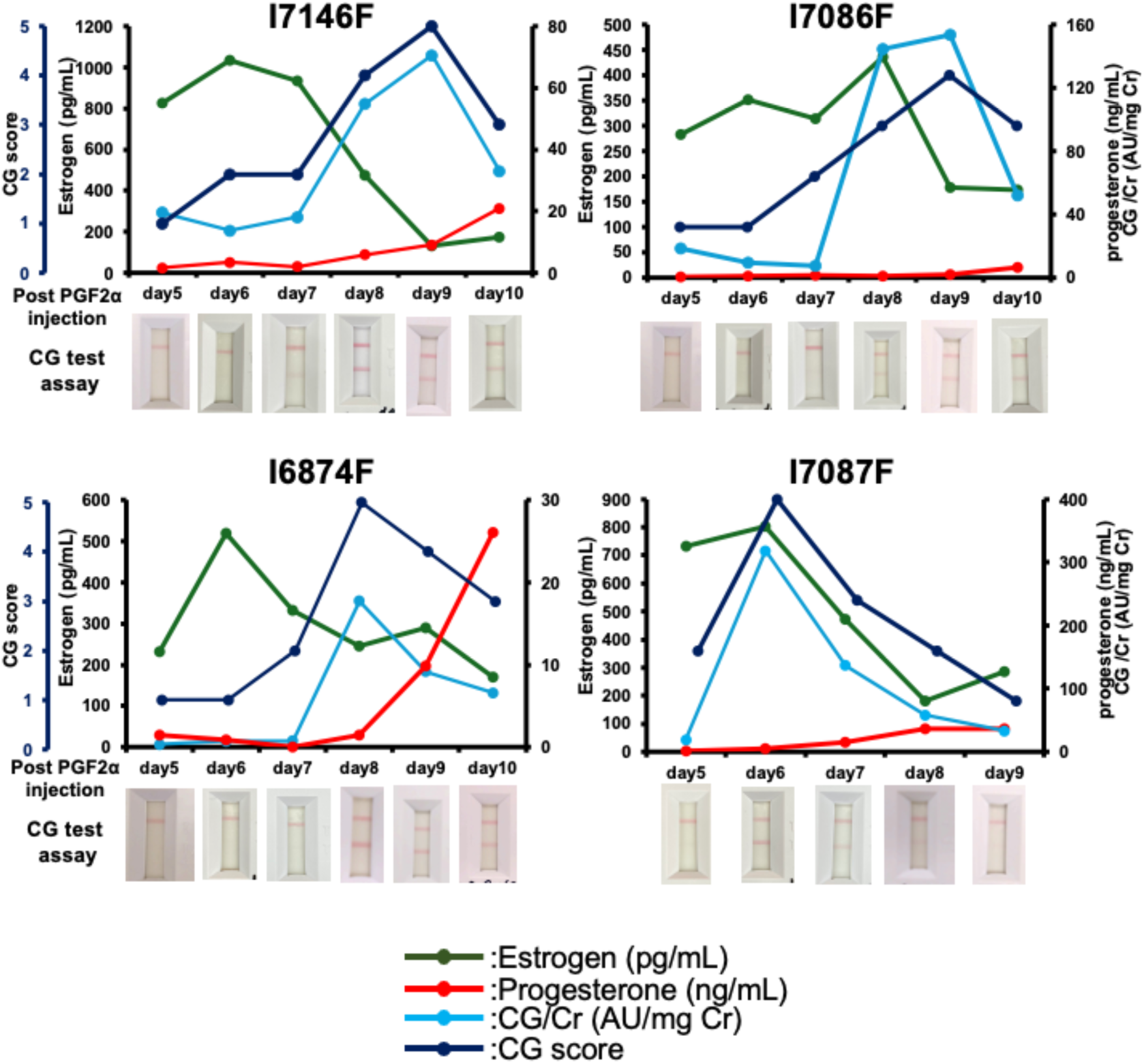
Detection of the CG surge before ovulation. A Sandwich ELISA analysis of a female marmoset’s urine was observed around ovulation. The marmoset IDs are I7146F, I7086F, I6874F, and I7087F. Green lines indicate plasma estradiol (pg/mL), red lines indicate plasma progesterone (ng/mL), light blue lines indicate urine CG (AU/mg Cr), and blue lines indicate CG score. The panels below the graph indicate the result of the CG assay. At the timing when a positive band (CG score is 3 or above) was detected, CG surge started. Moreover, the estradiol peak was after PGF2α injection day 6.5±1.15, and the CG peak was day 8.0±1.41. After the estradiol peak and CG peak was detected, progesterone was increased at day 9.5±1.25 after PGF2α injection.

### Success rate of artificial insemination based on CG surge before ovulation

To clarify whether the urinary CG positive band result of the CG assay prior to elevation in plasma P4 indicates ovulation, artificial insemination was performed on the day of or the day after the CG positive band was detected by the CG assay. Embryos were collected 5-8 days after artificial insemination from marmoset females via non-surgical uterine flushing. The results of embryo retrieval are shown in Table 2 and Supplementary Tables S2 and S3. The embryo recovery rate was 75% when artificial insemination was performed using the CG assay results as an indicator of ovulation. This outcome was significantly higher (p<0.05) than the 33.3% embryo recovery rate observed when artificial insemination was performed using the plasma P4 levels at a concentration exceeding 10 ng/ml (Table 2 and Supplementary Table S3). Furthermore, when artificial insemination was performed based on the results of the CG assay, outcomes of embryo retrieval were 100%, including dead embryos and nonfertilized oocytes. The resulting embryo retrieval rates indicate that the CG assay reflects the timing of ovulation. Additionally, embryos obtained 6–7 days after a positive CG assay were at the morula to compaction morula stage, indicating accurate ovulation timing.

**Table 2.** The success rate of AI.

| Hormone | No. of AI | Fertilized embryo (%) | Nonfertilized embryo (%) | Dead embryo (%) | Nothing (%) |
| --- | --- | --- | --- | --- | --- |
| Chorionic Gonadtropin | 12 | 9 (75.0)* | 1 (8.3) | 2 (16.7) | 0 |
| Progesterone | 12 | 4 (33.3) | 5 (41.7) | 0 | 3 (25.0) |

\*p<0.05 (X<sup>2</sup>-test)

Furthermore, in collaboration with another research institute, we also determined whether naturally mated embryos could be obtained by monitoring ovulation with a CG assay. Of the total 53 attempts, fertilized or unfertilized eggs, including dead embryos, were obtained in 60.4% of cases (Supplementary Table S1).

### Detection of CG during pregnancy

Female marmosets typically ovulate and become pregnant at approximately 10 days after delivery. To investigate the relationship between their reproductive cycle, including pregnancy and CG levels, CG levels were measured from the day after birth and monitored until their next pregnancy (Fig. 3, Table 3, and Supplementary Table S4). As a result, urine CG levels increased on average 10.67 ± 1.75 days after delivery, remained detectable for 2.3 ± 1.4 days, and then decreased below the detection level of the CG assay. Further, the first positive detection preceded the rise in plasma P4 levels, suggesting ovulation and day 0 for pregnancy. A second CG surge was observed 28.0 ± 2.4 days after delivery (17.3 ± 1.86 days after the first CG surge), persisting for 12.5 ± 1.09 weeks after the CG detection of the last delivery. Finally, the marmoset newborns were delivered 145.17 ± 1.47 days from the first day of the CG detection.

**Fig. 3.**
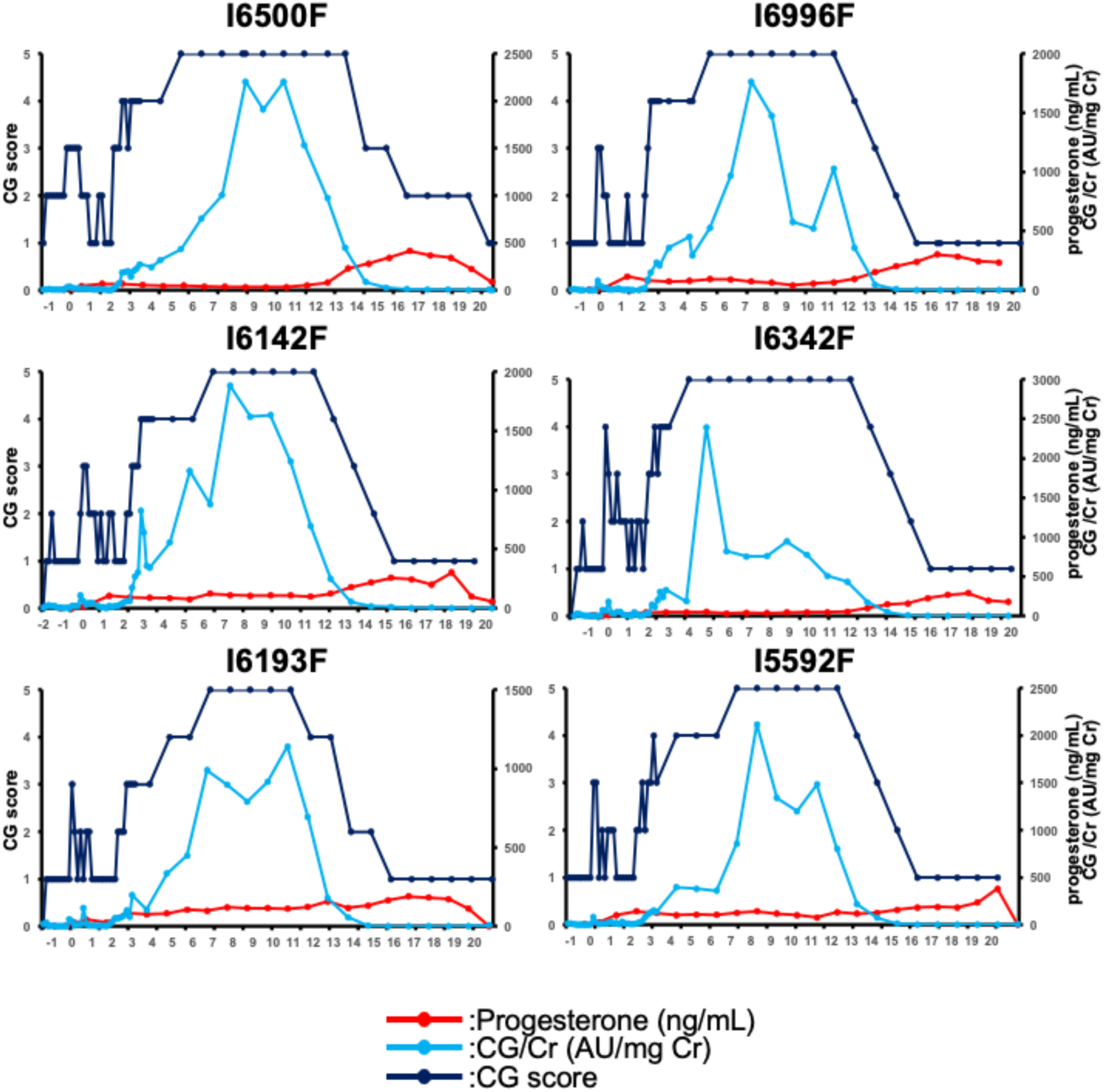
Urine CG transition from a marmoset birth to the next marmoset birth. A Sandwich ELISA analysis of a female marmoset’s urine was observed during pregnancy. The marmoset IDs are I6500F, I6996F, I6142F, I6342F, I6193F, and I5592F. Red lines indicate plasma progesterone (ng/mL), light blue lines indicate urine CG (AU/mg Cr), and blue lines indicate CG score. Each period is summarized on Table 3.

**Table 3.**
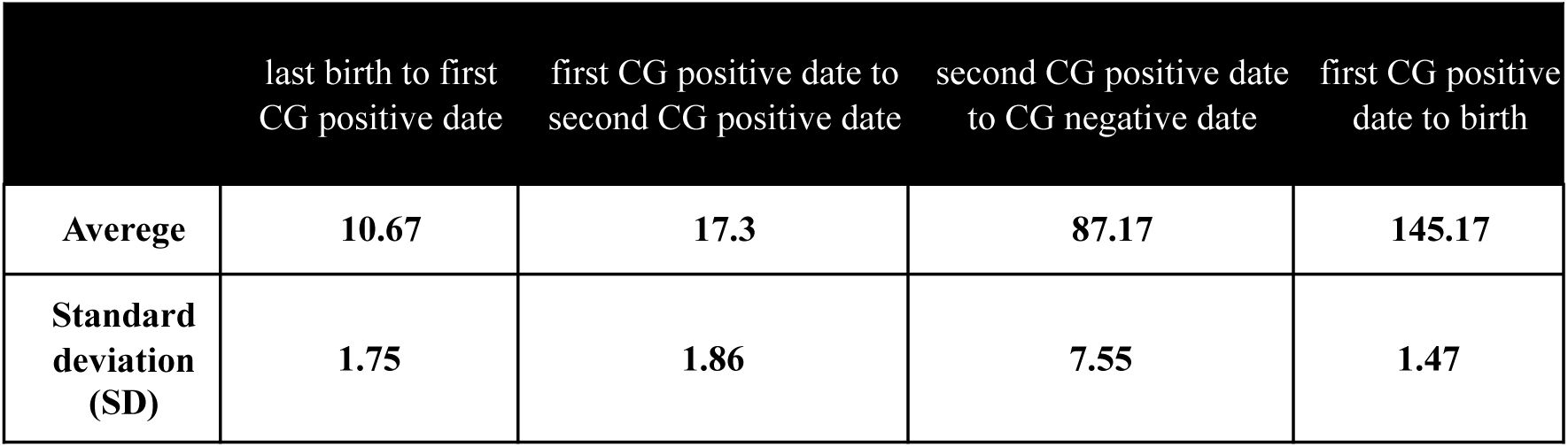
Transition in CG concentration and the sexual cycle.

|  | last birth to first CG positive date | first CG positive date to second CG positive date | second CG positive date to CG negative date | first CG positive date to birth |
| --- | --- | --- | --- | --- |
| Average | 10.67 | 17.3 | 87.17 | 145.17 |
| Standard deviation (SD) | 1.75 | 1.86 | 7.55 | 1.47 |

In contrast, ultrasound diagnoses showed that implantation could be confirmed at 29.6 ± 1.75 days after delivery and 17.3 ± 1.86 days after the first CG detection by the CG assay. All animals showing the second CG secretions were diagnosed as pregnant. These results indicate that the CG assay can accurately diagnose pregnancy from day 17.3 ± 1.86 days after ovulation.

Furthermore, the concentrations and scores of urine CG of all the data indicated a strong correlation (R=0.71). These results suggest that the CG assay results reliably reflect the CG concentration in urine (Figs. 2 and 3).

### Detection of pregnancy in other New World monkeys

To evaluate whether the CG assay could be applied to other New World monkeys, pregnancy diagnosis was performed every 3 weeks using urine samples from 8 squirrel monkeys. All five squirrel monkeys that showed continuous CG secretion gave birth within 20 weeks (Table 4). In contrast, the CG assay failed to identify pregnancy status in cynomolgus monkeys, an Old World monkey; neither 5 pregnancies nor 5 non-pregnancies were detected (Supplementary Table S5).

**Table 4.**
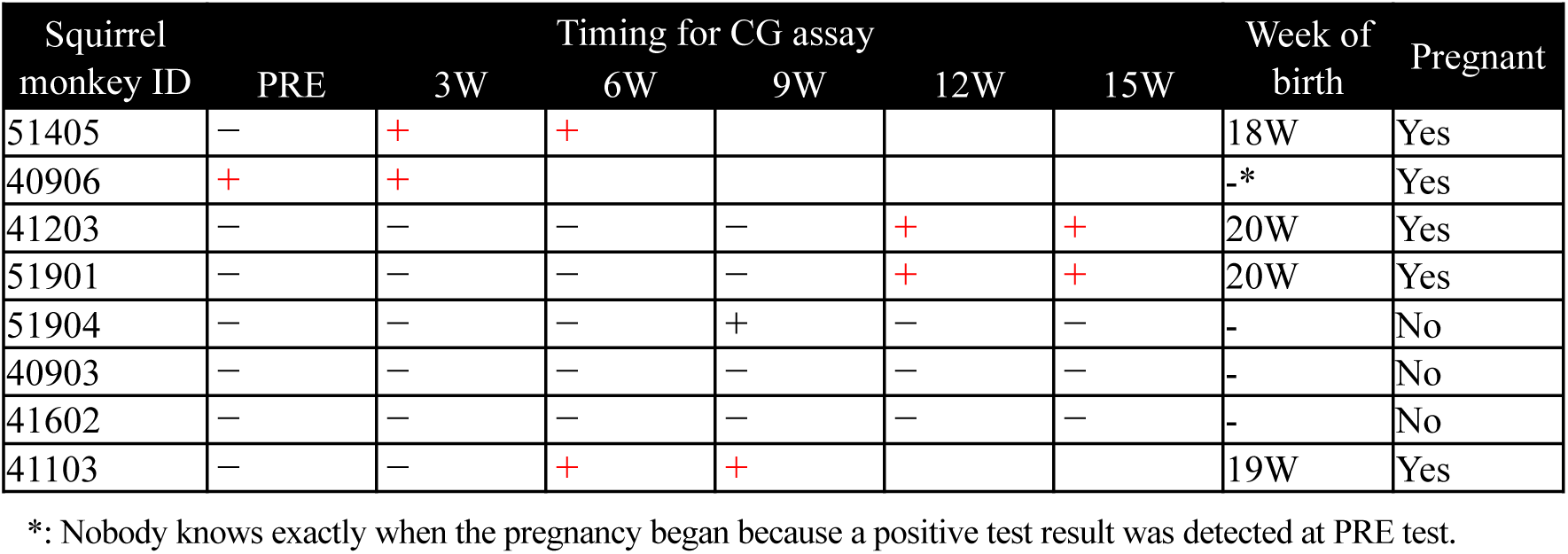
Pregnancy determination and results of birth in squirrel monkeys using the CG assay.

## Discussion

This study successfully developed novel anti-marmoset CG monoclonal antibodies and an immunochromatographic CG assay based on these antibodies. Although the assay was developed using marmoset CG, its successful application to squirrel monkeys suggests that it preferentially recognizes conserved CG features shared among certain New World primates rather than being strictly species restricted. The CG scores showed a strong correlation with ELISA measurements (R = 0.71), validating the reliability of this approach. To date, anti-human CG antibodies used in human pregnancy test kits fail to cross-react with marmoset CG, necessitating reliance on serial plasma progesterone measurements or ultrasonographic examination to confirm ovulation and pregnancy. In contrast, the CG assay developed in this study enables simple and early detection of ovulation and pregnancy in marmosets using non-invasive urine samples.

Consistent with previous reports, this study demonstrates that the plasma E2 surge preceded the urinary CG surge, followed by a subsequent rise in plasma P4 ^12^. However, the duration and timing of the E2 surge, gonadotropin peak, and P4 elevation observed in our study were shorter than those previously reported. Whereas earlier studies described E2 peaks occurring 7-9 days, gonadotropin peaks 9-10 days, and ovulation 9-14 days after PGF2α administration, our colony exhibited average E2 peaks at day 6.5, CG peaks at day 8.0, and P4 elevation at day 9.25. These differences may reflect variations in detection sensitivity associated with the measurement methods employed. Alternatively, they may be attributable to colony-specific characteristics, as our marmoset colony has been maintained as a closed population since 1983, potentially leading to genetic or environmental influences on reproductive endocrine dynamics. Importantly, despite these temporal differences, the relative sequence of endocrine events remained conserved, supporting the biological validity of CG as a marker for ovulation timing.

Three out of four animals showed a positive CG band in the CG assay one day after the initial decline in E2 levels, followed by a slight re-increase in E2 levels one to two days later. This initial decline in E2 coincided with the increase in urinary CG, consistent with the onset of ovulation. In the remaining case, E2 levels increased rapidly, making it difficult to define the precise day of decline. These endocrine patterns were consistent with previous reports ^13,14^. Artificial insemination performed immediately after CG elevation enabled the retrieval of fertilized embryos from the marmoset uterus several days later. In addition, naturally conceived embryos were successfully obtained by detecting ovulation using the CG assay. The embryo retrieval rate was significantly higher when ovulation was determined based on CG elevation than when based on P4 concentration (P < 0.05). Together, these findings indicate that urinary CG elevation precedes ovulation in marmosets and that the CG assay reliably captures the ovulatory window. Both artificial insemination and naturally conceived embryo retrieval outcomes demonstrate that ovulation can be detected using this simple, non-invasive method without the need for specialized equipment.

Moreover, very early pregnancies were detected using the CG assay at an average of 17.3 ± 1.86 days after ovulation, frequently preceding confirmation by ultrasonographic examination, which typically allows pregnancy diagnosis only after approximately 20 days post-ovulation. Longitudinal assessment of CG scores during pregnancy revealed that peak CG levels were maintained from approximately 5.5 to 11.3 weeks of gestation, declined to undetectable levels by around 14.5 weeks, and remained low thereafter. These CG secretion profiles are consistent with previous reports, suggesting that the CG assay captures CG derived from both the pituitary and the placental sources ^12^. Together, the detection of CG secretion across ovarian cycles and gestation supports the accuracy and biological relevance of the CG assay, which is further corroborated by successful embryo retrieval and subsequent live births of marmoset newborns.

Previous studies have strongly suggested that in marmosets, a CG surge occurs in place of the canonical LH surge during ovulation ^15^. However, the temporal relationship between CG elevation and ovulation has remained unclear. In this study, the CG assay enabled detection of both the peri-ovulatory period and early implantation, indicating that CG elevation reliably marks key reproductive transitions. These observations are consistent with the proposed model in which CG plays a central role in the ovulatory endocrine axis of New World primates. Nevertheless, definitive functional discrimination between CG- and LH-mediated signaling will require future studies employing antibodies or assays that selectively distinguish CG from LH.

The developed CG assay successfully detected pregnancies in squirrel monkeys (Table 4) but not in cynomolgus monkeys (Supplementary Table S5). The amino acid sequence identity of the CGα/β subunits between marmosets and squirrel monkeys (86.59%) was higher than between marmosets and the cynomolgus monkeys (73.33%), likely facilitating cross-reactivity of the anti-CG antibodies. This evolutionary conservation implies that our non-invasive monitoring platform could be broadly applicable to various Platyrrhini species, facilitating comparative reproductive studies without the physiological stress of blood sampling. Consistent with marmoset reproductive endocrinology, New World primates lack pituitary LH expression and instead rely on CG as the primary luteotropic hormone ^15–17^. The cross-reactivity of the developed CG assay in both marmosets and squirrel monkeys underscores the evolutionary conservation of the molecular architecture and functional domains of CG among New World primates.

These findings suggest that the CG assay preferentially recognizes conserved CG features shared among certain New World primates, rather than being universally applicable across primate lineages. Because reliable assessment of CG secretion dynamics during ovulatory cycles and pregnancy is essential for efficient breeding management, this assay may have potential utility in conservation-oriented reproductive monitoring programs, including those associated with initiatives such as the AZA Species Survival Plan (SSP). This possibility is further supported by the high amino acid sequence identity of CG among certain endangered New World primates, including cotton-top tamarins (95.73%), brown-headed tamarins (93.29%), and owl monkeys (86.49%), highlighting the broad utility of this assay for the advancement of conservation and reproductive management of diverse endangered New World primates.

In conclusion, the CG assay developed in this study provides a simple and efficient method for early pregnancy diagnosis and ovulation detection in marmosets. By enabling non-invasive monitoring of CG dynamics, this approach supports reproductive research in marmosets and offers a foundation for future extension to other New World primate species following appropriate validation. Furthermore, given the emerging evidence of significant differences in early embryonic development between rodents and primates, including humans, our assay provides a crucial platform to accelerate research using non-human primate embryos, ultimately bridging the gap in our understanding of early primate development.

## Materials and methods

### Animals

All animal experiments were approved by the Institutional Animal Care and Use Committee of the Central Institute for Experimental Medicine and Life Science (CIEM, approval nos. 20049A for 2022, AIA230050-001 for 2023, and AIA240014 for 2024 and 2025); CLEA Japan, Inc. (approval nos. 55-042, 56-026, 57-052 for mouse immunization and 57-015, 58-014, 59-036 for marmoset pregnancy diagnosis); National Institutes of Health/ National Institute of Neurological Disorders and Stroke (NIH/NINDS, approval nos. 1187 and 1196); Amami Laboratory of Medical Science, The Institute of Medical Science, The University of Tokyo (approval no. A2025IMS114-01); and Tsukuba Primate Research Center, National Institutes of Biomedical Innovation (approval no. DSR 07-32).

Six female common marmosets with normal ovarian cycles and four male common marmosets were used to obtain preimplantation embryos by artificial insemination at the CIEM. Six female common marmosets were used for pregnancy diagnosis at CLEA Japan, Inc. Six pairs of common marmosets with females of normal ovarian cycles and males were used to obtain preimplantation embryos by natural mating at the NIH/NINDS. All common marmosets were aged > 1.5 years. Moreover, eight squirrel monkeys were used for pregnancy diagnosis at the Amami Laboratory of Injurious Animals, Institute of Medical Science, University of Tokyo. All squirrel monkeys were aged > 4 years. Furthermore, cynomolgus monkeys were used for pregnancy diagnosis at Tsukuba Primate Research Center, National Institutes of Biomedical Innovation. They were aged > 5 years.

### Production of recombinant marmoset CG

Artificial DNA fragments encoding the translated regions of marmoset CGα (NCBI-Gene ID: 100389264) and CGβ (NCBI-Gene ID: 100391685) were synthesized (Eurofins Genomics, Tokyo, Japan). CGα cDNA was cloned into a pMT2 vector (Genetics Institute, MA, USA) with a C-terminal FLAG tag (CGα-FLAG), while CGβ cDNA was cloned into a pSecTag2 vector (Thermo Fisher Scientific, MA, USA) with an N-terminal PA tag (CGβ-PA). The CGα and CGβ expression constructs were transfected into CHO (ECACC 94060607) and CHO-S cells (Thermo Fisher Scientific), respectively, and these stably transfected cell lines were selected by αMEM (WAKO, Osaka, Japan) with MTX (WAKO), 10% fetal bovine serum (Corning), and Zeocin (Thermo Fisher Scientific). To obtain a stable CHO cell line producing both CGα and CGβ (recombinant CG) in a cell, the CGβ expression construct was transfected into the CGα expression cell line, and the stably transfected cell line was selected with Zeocin. These cell culture supernatants were subject to SDS-PAGE and Western blotting to identify the expression. CGα-FLAG and CGβ-PA proteins were purified from the cell culture supernatants using anti-DDDDK tag beads (MBL, Tokyo, Japan) and anti-PA tag beads (WAKO) per the manufacturer’s protocol. As shown in Supplementary Fig. S1A, the recombinant successful production of CG, comprising FLAG-tag fused CGα-FLAG and PA-tag fused CGβ-PA, was confirmed.

### Production of anti-CGα and CGβ monoclonal antibodies

Six-week-old female BALB/cA mice (CLEA Japan, Inc., Tokyo, Japan) were immunized by injections of purified CGα-FLAG or CGβ-PA proteins, and the hybridomas were generated from their lymphocytes, which were harvested from spleen and lymph node by fusing myeloma cells (P3U1) using a standard protocol. The obtained hybridomas were screened by ELISA using the supernatant and CGα-FLAG or CGβ-PA, and recombinant-CG as antigens. Fourteen hybridomas were selected from the anti-CGα antibody, and eight hybridomas were selected from the anti-CGβ antibody (Supplementary Table S6).

The class and subclass of the selected clones were determined by a mouse isotyping kit (Roche, North Ryde, Australia). The selected clones were cultured in Hybridoma-SFM (Gibco, NY, USA), and each antibody was purified using a Protein G Sepharose column (Cytiva, Uppsala, Sweden). Immunoprecipitation using recombinant-CG protein and the mAbs, and detection by Western blotting using the CGα -FLAG and CGβ-PA, confirmed that all mAbs except No. 1 shown in Supplementary Table S6 recognized recombinant-CG.

### Immunoprecipitation (IP)

IP was performed using the culture supernatant of recombinant CG as the antigen and the 14 anti-CGα mAb and 8 anti-CGβ mAb clones as the antibody. The collected samples were subjected to detection by Western blotting using anti-PA tag and anti-FLAG antibodies, respectively. The conditions for Immunoprecipitation and Western blotting were as follows. For Immunoprecipitation, 5 μg of each antibody was added to 50 μL of Sure Beads (Bio-Rad), and the culture supernatant of CG was reacted for 2 h at room temperature. The electrophoresis was performed using SuperSep™ Ace (12.5%, WAKO) at 200 V for 40 min under reducing conditions. Transcription was performed using Polyvinylidene Difluoride Membrane (Hydrophobic 0.2 μm, Thermo Fisher Scientific) at 20 V for 1 h. Immunoprecipitation samples with anti-CGα mAb and anti-CGβ mAb were detected using anti-PA tag antibody (WAKO) and anti-FLAG antibody (Abcam) (Supplementary Figs S1B and S1C).

### Sandwich ELISA for the detection of urinary marmoset CG

Each purified anti-CGα monoclonal antibody was conjugated to horseradish peroxidase (HRP) using HRP-labeling kits (DOJINDO, Kumamoto, Japan). The purified anti-CGβ and anti-CGα conjugated with HRP monoclonal antibodies were used as the capture and detection antibodies to determine the optimal combination. Sandwich ELISA was performed using microtiter plates; the procedure was performed as follows: MaxiSorp 96-well plates (#442404, Thermo Fisher Scientific) were coated with 50 µL of the anti-CGβ monoclonal antibody (5 µg/mL) in phosphate-buffered saline (PBS) and incubated overnight at 4℃. After washing with PBS containing 0.05% Tween20 (PBS-T) once, the well surfaces were blocked with 200-300 µL of blocking reagent (#IS-CD-500E, Cosmo Bio Co., Ltd.) for 3-5 h at room temperature. After removing the solution and drying up the plate at room temperature completely, 50 µL of two-fold serially diluted standards (the cell culture supernatants of CGα and CGβ expression CHO cell line) by ten-fold diluted Blocking one (Nacalai Tesque inc., Kyoto, Japan) as a dilution buffer, and urine specimens were added, and incubated at room temperature for 1 h. After washing with PBS-T three times, the 50 µL anti-CGα conjugated with HRP monoclonal antibody (1: 10000-30000 in dilution buffer) was added and incubated at room temperature for 1 h. After washing with PBS-T four times, 50 µL of substrate (TMB, Thermo Fisher Scientific) was added and incubated at room temperature for 20 min. The reaction was stopped by 50 µL of 1-2N sulfuric acid. Absorbance was measured at 450 nm by the microplate reader (Multiskan FC, Thermo Fisher Scientific). The ELISA standard was collected from stable CG expressing CHO cells cultured in serum-free αMEM and the concentration was determined as 1,000 arbitrary units per milliliter (AU/mL). The urinary CG concentrations of each sample were calculated based on the standard curve obtained and the measured concentrations were adjusted by those of urinary creatinine (AU/mg Cr).

### Development of the CG assay

Anti-mouse immunoglobulin and anti-CGα monoclonal antibodies were immobilized onto a nitrocellulose membrane for the control and test lines as capture antibodies. A reagent pad containing anti-CGβ monoclonal antibody conjugated with colloidal gold was located between the test lines and the sample well. The assay was initiated by adding 90 μL of the test solution to the sample well. The membrane was observed visually after a 5-min incubation period at room temperature. A red line in the control line indicates the absence of the marmoset CG. The concurrent presence of red lines in both the control and test lines indicates the presence of marmoset CG.

The best pair for anti-CGα mAb and anti-CGβ mAb was selected by sandwich ELISA. Eight anti-CGβ mAbs were captured, and five anti-CGα mAb (5H3, 46A4, 52B7, 59E1, and 65C10; Supplementary Table S7), representing the top five for high activity, were used for detection. Forty pairs of anti-CGα mAbs and anti-CGβ mAbs were tested using sandwich ELISA to identify the best combination for immunochromatography (Supplementary Table S7). Eight anti-CGβ mAbs were evaluated alongside the top five highly active anti-CGα mAbs used for detection. A pair of anti-CGα mAb No.5 (clone# 52B7) for capturing antibodies and anti-CGβ mAb No.18 (clone# 13B6) for labeling antibodies, shown in Supplementary Table S7, was used for the development of the CG assay (Supplementary Figs S2A and S2B). A CG score of “3” was assigned when the test line (lower band) was detected within 5 min.

### Pregnancy diagnosis

Marmosets’ pregnancies were confirmed using the ultrasonographic examination (uSmart 3200T, Terason, MA, USA) and the developed CG assay. When the endometrium was closed and the CG assay was negative, the animals were considered nonpregnant, whereas when the endometrium was separated and a lumen was visible and the CG assay was positive, the animals were considered pregnant. Moreover, after 40 days of pregnancy, the diameter of the uterus was measured by uterine palpation under no anesthesia to confirm pregnancy status.

Squirrel monkeys’ pregnancies were checked every 3 weeks during breeding season. The animals that showed a positive band in the CG were evaluated based on whether they gave birth at approximately 20 weeks later. Cynomolgus monkeys’ pregnancies were confirmed by ultrasonographic examination (ARIETTA 750SE, Fujifilm Corporation, Tokyo, Japan) and measurement of plasma P4 and E2 levels.

### Preparation of sperm

Fresh semen was collected by the Penile Vibratory Stimulations method, with minor modifications from previous protocols ^18^. Briefly, a handy massager was used as the vibrating shaft, and a 200 μL pipette tip and silicone tubing (about 30 mm; internal diameter, 3.3 mm; outer diameter, 5 mm) were thrust into the silicone surface of the massager. Unsedated males were restrained in a marmoset restrainer (CL-4532; CLEA Japan, Inc.), and both legs were held manually. The penis was gently extruded from the preputial sheath, and the vibrating silicon tube was held against the preputial orifice.

Semen samples were diluted with 200 μL TYH medium (LSI Medience Corporation, Tokyo, Japan) prepared in a conical tube at room temperature. The samples were gently mixed using a 1000 μL pipette and centrifuged at 250 g for 5 min. The supernatant was discarded, and 100 μL of fresh TYH medium was gently mixed using a 1000 μL pipette. Sperm concentration and motility were determined using SMAS (DITECT, Tokyo, Japan, Supplemental Tables S2 and S3). Depending on the semen volume, sperm concentrations were adjusted to 1.56 to 21.13 X 10^6^/μL (mean: 11.13 ± 5.61 × 10^6^/μL), and sperm motilities ranged from 37.07 % to 83.38 % (mean: 64.59 ± 15.25%).

### Detecting ovulation

Four female marmosets with regular ovulation cycles were administered PGF2α analog cloprostenol (Estrumate; Nagase Medicals Co., Ltd, Hyogo, Japan) more than 10 days after the beginning of the luteal phase as previously described ^19^. Urine samples and plasma samples were collected from day 5 to day 10 after PGF2α injection. Urine samples were collected from the aluminum floorboards of the isolation box by isolating the female for up to 4h in the isolation box, which allowed for visual and auditory contact of the female with cage mates.

The ovulation was monitored by measuring the levels of urine CG using the CG assay and plasma P4 and E2 using an EIA kit (TOSOH, Tokyo, Japan). Ovulation was determined as the day before serum progesterone levels exceeded 10 ng/mL, and the detection of E2 and CG peaks.

### Artificial insemination

Six adult female marmosets with regular ovarian cycles were used for artificial insemination. Ovarian cycles were controlled by the PGF2α analog cloprostenol (Estrumate; Nagase Medicals Co., Ltd, Hyogo, Japan), as described above. The ovarian cycles were monitored by measuring the plasma P4 concentration using an EIA kit (TOSOH, Tokyo, Japan) or the levels of urinary CG using the CG assay. Ovulation was determined as a day before serum progesterone levels exceeded 10 ng/mL or a day after urine CG detection by the CG assay.

Artificial insemination was performed on ovulation day or the next day. A 100 μL aliquot of diluted fresh sperm was injected into the female marmoset’s vagina using a sterile plastic feeding catheter for mice (length: 25 mm; inner diameter: 0.68 mm; outer diameter: 1.18 mm; and first ball diameter: 2 mm; Fuchigami, Kyoto, Japan) attached to a 1 mL syringe (TERUMO, Tokyo, Japan).

### Collection of preimplantation embryos

Preimplantation embryos were collected as previously described ^20,21^. The preimplantation embryos were obtained via nonsurgical embryo collection, with embryos from the 8-cell to blastocyst stages, after ovulation on days 5-8, acquired using this procedure. The anesthesia was performed using the same protocol as previously reported, and all devices used for embryo collection were identical to those for embryo transfer ^22^. Female marmosets were pre-anesthetized using 0.04 mg/kg medetomidine (Domitor; Nippon Zenyaku Kogyo, Fukushima, Japan), 0.40 mg/kg midazolam (Dormicam; Astellas Pharma, Tokyo, Japan), and 0.40 mg/kg butorphanol (Vetorphale; Meiji Seika Pharma, Tokyo, Japan). During the operation, the marmosets were anesthetized via isoflurane (DS Pharma, Osaka, Japan) inhalation and kept on warm pads. A blunt-ended stainless-steel stylet (25G; 120-mm long) covered with the 18G, 108-mm long Fluon ETFE catheter (Oviraptor; Altair, Kanagawa, Japan) was inserted into the uterus through the uterine cervix. The blunt-end stainless steel stylet was then removed, and insertion of the outer Fluon ETFE catheter was retained for flushing. The 5 ml syringe was connected to an extension tube (1.0 mm in diameter, 40 cm long) attached to a polyethylene cannula (160 mm long; inner diameter: 0.28 mm; and outer diameter: 0.61 mm; Altair), which was filled with flushing medium [Waymouth’s medium (Thermo Fisher Scientific) and 1 M HEPES (Sigma-Aldrich, MO, USA)]. The catheter with flushing medium was then inserted into the uterus through the polyethylene cannula, and the medium was flushed into the uterus. During flushing, the oviducts were compressed against the uterus by palpation through the abdominal wall to prevent the embryos from being flushed out through the fimbria ovarica. The perfused medium was administered through a Fluon ETFE catheter and collected in a Petri dish to retrieve fertilized preimplantation embryos. All procedures were performed under sterile conditions.

## Author contributions

K.K., T.S., K.F., and E.S. designed the study. K.K., T.S., and E.S. designed and analyzed the data. A.I., M.H., T.S., Y.B., and K.F. made new anti-CG Abs and developed the CG assay. S.K. and T.M. performed the CG assay for the squirrel monkey. M.K. and T.S. performed the CG assay for the cynomolgus monkey. H.N., J.B., S.O., M.H., L.B., W.K., Y.K., Y.Y., K.K., and E.S. performed the embryo collection at NIH. M.K., J.A., and Y.Y. contributed technical support, reagents, and materials. K.K., T.S., A.I., and M.H. performed the experiments and acquired the data. K.K., T.S., and E.S. interpreted the data and wrote the manuscript. E.S. obtained financial support.

## Acknowledgments

We would like to thank Takashi Inoue, Keisuke Mukasa, Terumi Yurimoto, Noriko Kato, Yoshimitu Togashi, Mai Aoyama, and all the members of Division of Advanced Physiology. We would also like to thank Editage (www.editage.com) for English language editing. This research was supported in part by the Intramural Research Program of the National Institutes of Health (NIH), National Institute of Neurological Disorders and Stroke (NINDS). The contributions of the NIH author(s) were made as part of their official duties as NIH federal employees, are in compliance with agency policy requirements, and are considered Works of the United States Government. However, the findings and conclusions presented in this paper are those of the author(s) and do not necessarily reflect the views of the NIH or the U.S. Department of Health and Human Services.

## Data availability statement

The authors confirm that the data supporting the findings of this study are available within the article and its supplementary materials.

