## Supplementary material for "A chorionic gonadotropin assay enables non-invasive detection of ovulation and early pregnancy in a New World primate model": Suplemental tables and figures

**Development of a species-specific chorionic gonadotropin immunochromatographic test kit for early pregnancy diagnosis and ovulation detection in common marmosets**

Keiko Kishimoto[1**], Takuma Soga[2**], Akio Iio[2,3], Masahiko Hatakeyama[2], Satoru Kawai[4], Michiko Kamioka[1], Jinsho Aoki[2], Yuka Bunzui[2], Yuko Yamada[1], Miho Kohara[5], Yoko Kurotaki[6], Wakako Kumita[1], Julie Brent-Cummins[7], Sang Su Oh[7], Milton Herrera[7], Lian Bik[7], Heather Narver[8], Tadashi Sankai[5], Tomoji Mashimo [4, 9], Kazumasa Fukasawa[2], and Erika Sasaki[1]*

**Supplementary Fig. S1: Development of novel marmoset anti-CG mAbs**

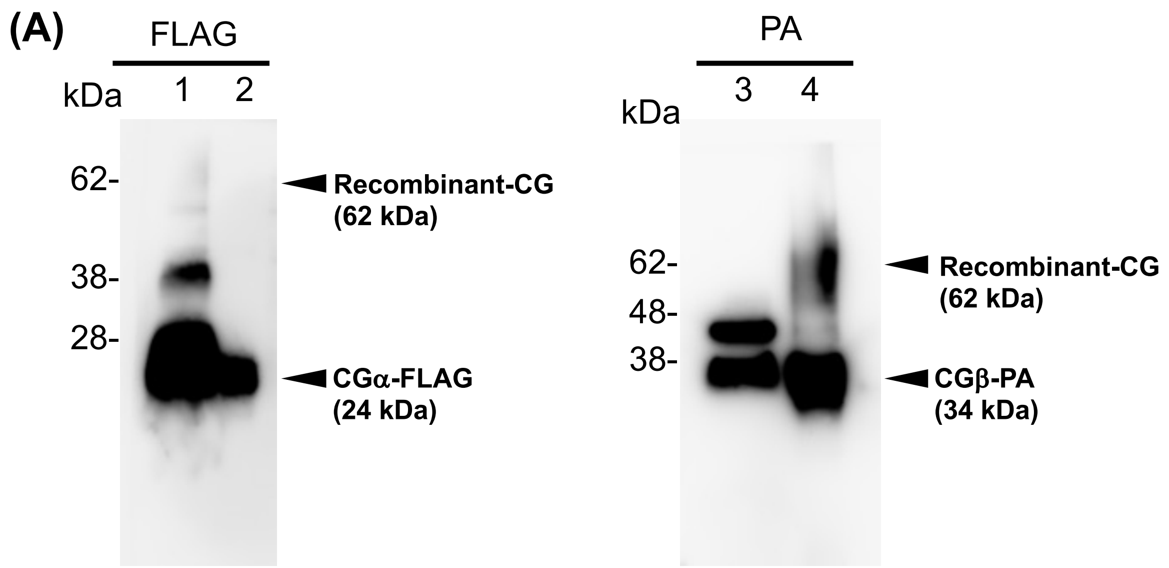


(A) Western blot detected recombinant protein expression. Lane 1: Supernatant of CHO cells expressed recombinant-CG purified by FLAG column and lane 2: Supernatant of CHO cells expressed CGα-FLAG purified by FLAG column were detected by anti-FLAG antibody. Lane 3: Supernatant of CHO cells expressed CGβ-PA purified by PA column and lane 4: Supernatant of CHO cells expressed recombinant-CG purified by PA column were detected by anti-PA antibody. CG: chorionic gonadotropin

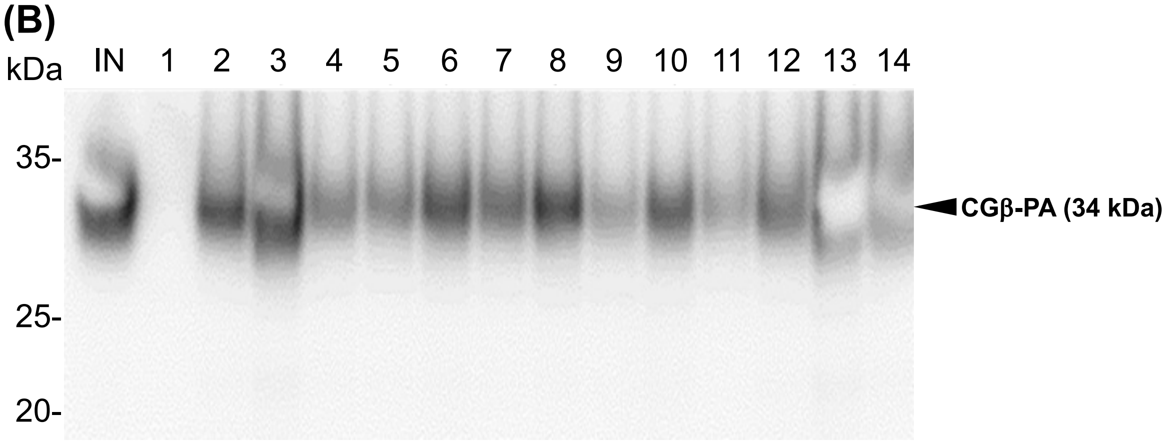


(B) Immunoprecipitation was performed using the novel marmoset anti-CGα mAbs obtained for the antibody and recombinant-CG protein for the antigen. Lane no. means antibody no. in Supplementary table S6 and is used for immunoprecipitation. IN means Input; supernatant of CHO cells expressing recombinant-CG. The detection used an anti-FLAG antibody recognized CGα-FLAG. CGβ-PA protein was detected at 34 kDa. CG, chorionic gonadotropin

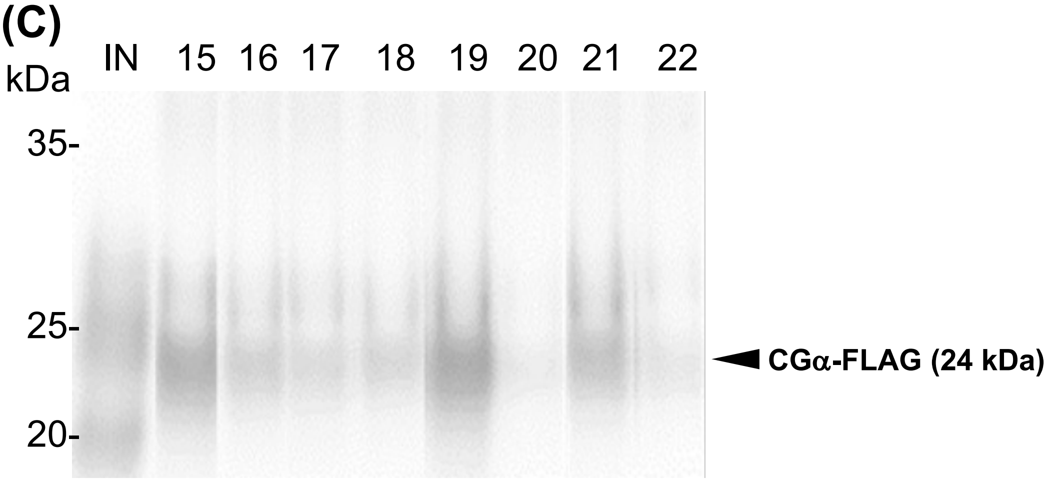


(C) Immunoprecipitation was performed using the novel marmoset anti-CGβ mAbs obtained for the antibody and recombinant-CG protein for the antigen. Lane no. means antibody no. in Supplementary table S6 and is used for IP. IN means Input; supernatant of CHO cells expressed recombinant-CG. The detection used an anti-PA antibody recognized CGβ-PA.

CGα-FLAG recombinant protein was detected at 24 kDa.

**Supplementary Fig. S2: The best combination for sandwich ELISA**

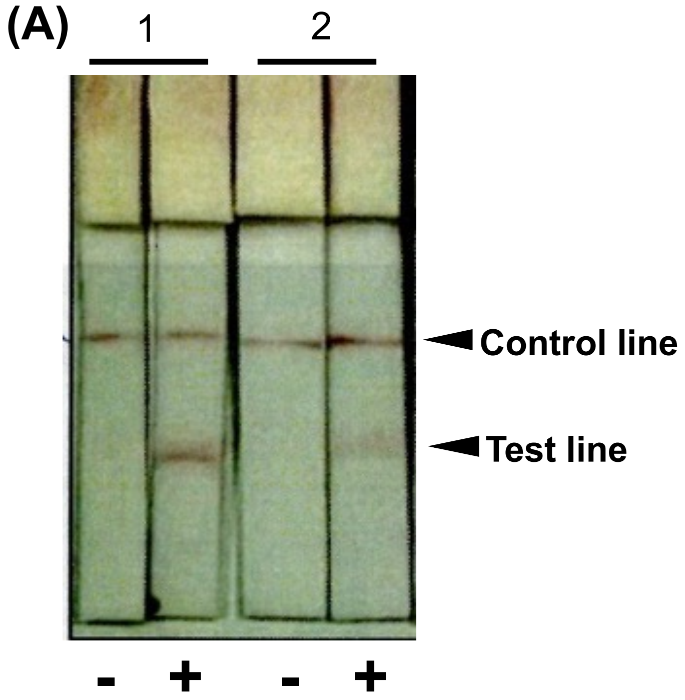


(A) The immunoassay was performed to develop the immunochromatographic test kit. Lane 1 is the labeled antibody 52B7 (#5 in Supplementary table S6), and the captured antibody is 13B6 (#18 in Supplementary table S6). Lane 2 is the labeled antibody 13B6 (#18 in Supplementary table S6), and the captured antibody is 52B7 (#5 in Supplementary table S6). "-" means the urine from no pregnant female marmoset, and "+" means the urine from the pregnant female marmoset. Although the test line detected both conditions, lane 1 is better than lane 2.

**(B)**

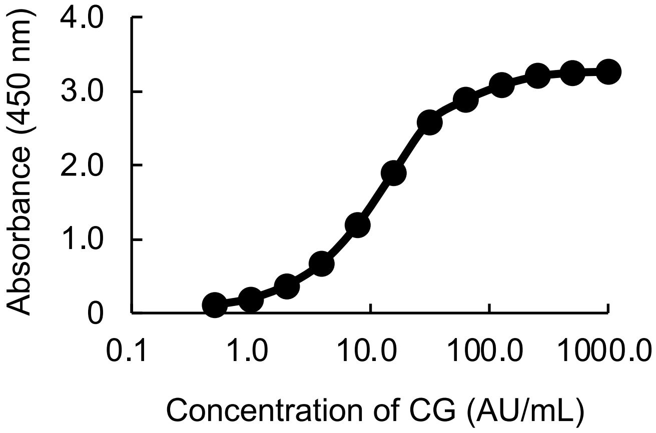


(B) The result of the sandwich ELISA shows that 52B7 was the labeled antibody, and 13B6 was the captured antibody. The graph indicates the relationship between the concentration of CG and Absorbance. When the concentration of CG was 100 AU/mL, Absorbance was plateaued.

**Supplementary table S1: The result of embryo collection at NIH**

| No. of embryo collection | Fertilized embryo (%) | Nonfertilized embryo (%) | Dead embryo (%) | Nothing (%) |
|---|---|---|---|---|
| 53 | 26 (49.1) | 2 (3.8) | 4 (7.5) | 21 (39.6) |

#### Supplementary table S2: The ovulation cycle measured by CG and timing of AI

| Females | | | Males | | | Result of AI | |
|---|---|---|---|---|---|---|---|
| ID (age) | Days after PGF2α injection | CG test kit | ID (age) | Sperm Concentration ($X10^6$/mL) | Motility (%) | embryo stage | Number of embryos |
| I7087F (2.0) | 7 | Negative | | | | | |
| | 8 | Positive | SI784tgM (8.8) | 21.13 | 83.38 | | |
| | 15 | | | | | EB, CM | 2 |
| I7146F(1.9) | 6 | Negative | | | | | |
| | 7 | Negative | | | | | |
| | 8 | Negative | | | | | |
| | 9 | Positive | I813tgM(7.4) | 20.38 | 76.9 | | |
| | 15 | | | | | N.F | 1 |
| I7087F (2.0) | 6 | Negative | | | | | |
| | 7 | Positive | I813tgM(7.5) | 3.67 | 78.94 | | |
| | 15 | | | | | CM, N.F | 2 |
| I7086F(2.0) | 6 | Negative | | | | | |
| | 7 | Negative | | | | | |
| | 8 | Positive | SIH/SI948tgM(2.9) | 4.92 | 61.47 | | |
| | 15 | | | | | CM, M | 2 |
| I7087F (2.1) | 6 | Negative | | | | | |
| | 7 | Negative | | | | | |
| | 8 | Positive | | | | | |
| | 9 | | I946tgM(2.1) | 8.80 | 73.85 | | |
| | 15 | | | | | 8C, 8C, M | 3 |
| I6874F (2.9) | 6 | Negative | | | | | |
| | 7 | Negative | | | | | |
| | 8 | Negative | | | | | |
| | 9 | Positive | I946tgM(2.1) | 8.80 | 73.85 | | |
| | 15 | | | | | CM, CM | 2 |
| I7086F(2.1) | 6 | Negative | | | | | |
| | 7 | Positive | I813tgM(7.5) | 9.03 | 37.07 | | |
| | 15 | | | | | dead | 1 |
| I7146F(1.9) | 6 | Negative | | | | | |
| | 7 | Negative | | | | | |
| | 8 | Negative | | | | | |
| | 9 | Positive | SIH/SI948tgM(2.9) | 10.85 | 51.69 | | |
| | 15 | | | | | 12C | 1 |
| I7002F(2.5) | 6 | Negative | | | | | |
| | 7 | Negative | | | | | |
| | 8 | Negative | | | | | |
| | 9 | Positive | I784tgM(8.9) | 16.21 | 40.89 | | |
| | 15 | | | | | M, M, M | 3 |
| I6934F(2.9) | 6 | Negative | | | | | |
| | 7 | Negative | | | | | |
| | 8 | Negative | | | | | |
| | 9 | Positive | I946tgM(2.1) | 13.42 | 75.18 | | |
| | 15 | | | | | M | 1 |
| I6934F(3.0) | 6 | Negative | | | | | |
| | 7 | Negative | | | | | |
| | 8 | Negative | | | | | |
| | 9 | Negative | | | | | |
| | 10 | Positive | I946tgM(3.0) | 6.983 | 56.38 | | |
| | 13 | | | | | M, dead | 2 |
| I7087F (2.5) | 7 | Negative | | | | | |
| | 8 | Positive | I946tgM(3.1) | 9.393 | 65.44 | | |
| | 15 | | | | | dead, N.F. | 2 |

### Supplementary table S3: The ovulation cycle measured by progesterone and timing of AI

| Females | | | Males | | | Result of AI | |
|---|---|---|---|---|---|---|---|
| ID (age) | Days after PGF2α injection | Progesterone (ng/mL) | ID (age) | Sperm Concentration ($X10^6$/mL) | Motility (%) | embryo stage | Number of embryos |
| I6874F(2.3) | 1 | 3 | | | | | |
| | 9 | 0.9 | SI784tgM (8.4) | 8.771 | 70.77 | | |
| | 12 | 21.8 | SI784tgM (8.4) | 9.084 | 64.46 | | |
| | 15 | | | | | M, CM, Bl | 3 |
| I6934F (2.1) | 1 | 4.2 | | | | | |
| | 9 | 4.2 | SI784tgM (8.4) | 5.944 | 63.97 | | |
| | 12 | 60.9 | | | | | |
| | 15 | | | | | N.F., N.F. | 2 |
| I6934F (2.2) | 1 | 4.1 | | | | | |
| | 7 | 1.5 | I813tgM (7.0) | 19.346 | 74.66 | | |
| | 9 | 3 | SIH/SI948tgM(2.4) | 12.112 | 71.82 | | |
| | 12 | 50.4 | | | | | |
| | 15 | | | | | M, N.F. | 2 |
| I6934F (2.2) | 1 | 2.2 | | | | | |
| | 7 | 1.8 | SIH/SI948tgM(2.5) | 11.036 | 75.22 | | |
| | 9 | 1 | SIH/SI948tgM(2.5) | 4.226 | 62.19 | | |
| | 12 | 48 | | | | | |
| | 15 | | | | | 6C, CM, dead | 3 |
| I6934F (2.3) | 1 | 2.4 | | | | | |
| | 7 | 2.2 | I946tgM(2.7) | 4.924 | 76.77 | | |
| | 9 | 1.5 | I946tgM(2.7) | 16.812 | 83.86 | | |
| | 12 | 23.4 | | | | | |
| | 15 | | | | | 12C, 12C, 12C | 3 |
| I6874F (2.7) | 1 | 1.8 | | | | | |
| | 6 | 1.8 | SIH/SI948tgM(2.6) | 1.560 | 82.40 | | |
| | 8 | 1.5 | SI784tgM (8.5) | 9.381 | 87.12 | | |
| | 12 | 51.8 | | | | | |
| | 14 | | | | | N.F. | 1 |
| I7087F (2.8) | 1 | 1.6 | | | | | |
| | 8 | 1.6 | SIH/SI948tgM(2.6) | 12.481 | 82.40 | | |
| | 10 | 9.8 | | | | | |
| | 16 | | | | | N.F. | 1 |
| I7087F (2.9) | 1 | 1.1 | | | | | |
| | 7 | 6.6 | I946tgM(2.8) | 12.528 | 87.50 | | |
| | 9 | 59.2 | | | | | |
| | 15 | | | | | N.F. | 1 |
| I6874F (2.8) | 1 | 4.3 | | | | | |
| | 7 | 0.8 | SI784tgM (8.6) | 8.669 | 37.56 | | |
| | 9 | 3.8 | SI784tgM (8.6) | 11.109 | 77.74 | | |
| | 11 | 49 | | | | | |
| | 17 | | | | | N.F. | 1 |
| I6934F (2.4) | 1 | 2.1 | | | | | |
| | 7 | 0.6 | I813tgM (7.1) | 6.370 | 65.33 | | |
| | 9 | 0.7 | SIH/SI948tgM(2.7) | 3.773 | 62.74 | | |
| | 12 | 17.9 | | | | | |
| | 15 | | | | | | 0 |
| I6934F (2.4) | 1 | 4.5 | | | | | |
| | 7 | 1.3 | SI784tgM (8.6) | 9.450 | 71.36 | | |
| | 9 | 1.3 | I813tgM (7.2) | 13.246 | 88.01 | | |
| | 13 | 20.4 | | | | | |
| | 15 | | | | | | 0 |
| I7002F (2.5) | 1 | 6 | | | | | |
| | 7 | 1.1 | I813tgM (7.2) | 12.803 | 77.30 | | |
| | 9 | 1.8 | I946tgM(2.9) | 11.546 | 83.57 | | |
| | 12 | 38.4 | | | | | |
| | 14 | | | | | | 0 |

### Supplementary table S4: The detail result for Fig.3

| I6500F | | | | | I6996F | | | | |
|---|---|---|---|---|---|---|---|---|---|
| Pregnancy date | *USG | **Progesterone (ng/mL) | Dual Checker Decision | Dual Checker Score | Pregnancy date | *USG | **Progesterone (ng/mL) | Dual Checker Decision | Dual Checker Score |
| Birth | | | | | Birth | | | | |
| 1 | | | − | 1 | 1 | | | − | 1 |
| 2 | | | − | 2 | 2 | | | − | 1 |
| 3 | | | − | 2 | 3 | | | − | 1 |
| 4 | | | − | 2 | 4 | 1.037 | | − | 1 |
| 5 | | | − | 2 | 5 | 1.038 | | − | 1 |
| 6 | | | − | 2 | 6 | 1.038? | | − | 1 |
| 7 | | 4.66 | − | 2 | 7 | 1.042 | 8.12 | − | 1 |
| 8 | | | − | 2 | 8 | 1.042 | | − | 1 |
| 9 | | | + | 3 | 9 | | | − | 1 |
| 10 | | 10.14 | + | 3 | 10 | 1.039 | 9.94 | + | 3 |
| 11 | | | + | 3 | 11 | 1.039 | | + | 3 |
| 12 | 1.028 | | + | 3 | 12 | 1.039 | 20.26 | − | 2 |
| 13 | 1.027 | 44.9 | + | 3 | 13 | 1.041 | | − | 2 |
| 14 | 1.027 | | − | 2 | 14 | 1.036 | | − | 1 |
| 15 | | | − | 2 | 15 | | | − | 1 |
| 16 | | | − | 2 | 16 | | | − | 1 |
| 17 | | | − | 1 | 17 | | | − | 1 |
| 18 | | | − | 1 | 18 | 1.033 | | − | 1 |
| 19 | | | − | 1 | 19 | 1.031 | | − | 1 |
| 20 | 1.026 | 68.42 | − | 2 | 20 | 1.035 | 115.8 | − | 2 |
| 21 | 1.028 | | − | 2 | 21 | 1.035 | | − | 1 |
| 22 | | | − | 1 | 22 | | | − | 1 |
| 23 | | | − | 1 | 23 | 1.034 | | − | 1 |
| 24 | | | − | 1 | 24 | 1.035 | | − | 1 |
| 25 | 1.029 | | + | 3 | 25 | 1.036 | | − | 1 |
| 26 | 1.027 | | + | 3 | 26 | 1.023 | | − | 2 |
| 27 | 1.02 | 66.84 | + | 3 | 27 | 1.036 | 81.04 | + | 3 |
| 28 | 1.029 | | + | 4 | 28 | 1.04 | | + | 4 |
| 29 | 1.028 | | + | 4 | 29 | | | + | 4 |
| 30 | | | + | 3 | 30 | 1.033 | | + | 4 |
| 31 | 1.015 | | + | 4 | 31 | 1.032 | | + | 4 |
| 32 | 1.027 | | + | 4 | 32 | | | | |
| 33 | 1.03 | | + | 4 | 33 | | | | |
| 34 | 1.024 | 55.92 | + | 4 | 34 | 1.04 | 72.88 | + | 4 |
| 35 | | | | | 35 | | | | |
| 36 | | | | | 36 | | | | |
| 37 | | | | | 37 | | | | |
| 38 | | | | | 38 | | | | |
| 39 | | | | | 39 | | | | |
| 40 | | | | | 40 | | | | |
| 41 | 1.025 | 46.2 | + | 4 | 41 | 1.045 | 78.12 | + | 4 |
| 42 | | | | | 42 | | | + | 4 |
| 43 | | | | | 43 | | | | |
| 44 | | | | | 44 | | | | |
| 45 | | | | | 45 | | | | |
| 46 | | | | | 46 | | | | |
| 47 | | | | | 47 | | | | |
| 48 | 1.03 | 48.2 | + | 5 | 48 | 1.036 | 94.52 | + | 5 |
| 49 | | | | | 49 | | | | |
| 50 | | | | | 50 | | | | |
| 51 | | | | | 51 | | | | |
| 52 | | | | | 52 | | | | |
| 53 | | | | | 53 | | | | |
| 54 | | | | | 54 | | | | |
| 55 | 1.027 | 40.92 | + | 5 | 55 | 1.038 | 90.2 | + | 5 |
| 56 | | | | | 56 | | | | |
| 57 | | | | | 57 | | | | |
| 58 | | | | | 58 | | | | |
| 59 | | | | | 59 | | | | |
| 60 | | | | | 60 | | | | |
| 61 | | | | | 61 | | | | |
| 62 | 1.01 | 36.24 | + | 5 | 62 | 1.038 | 74.36 | + | 5 |
| 63 | | | | | 63 | | | | |
| 64 | | | | | 64 | | | | |
| 65 | | | | | 65 | | | | |
| 66 | | | | | 66 | | | | |
| 67 | | | | | 67 | | | | |
| 68 | | | | | 68 | | | | |
| 69 | | 34.44 | + | 5 | 69 | 1.035 | 65.12 | + | 5 |
| 70 | | | | | 70 | | | | |
| 71 | | | | | 71 | | | | |
| 72 | | | | | 72 | | | | |
| 73 | | | | | 73 | | | | |
| 74 | | | | | 74 | | | | |
| 75 | | | | | 75 | | | | |
| 76 | 1.018 | 33.44 | + | 5 | 76 | 1.016 | 42.52 | + | 5 |
| 77 | | | | | 77 | | | | |
| 78 | | | | | 78 | | | | |
| 79 | | | | | 79 | | | | |
| 80 | | | | | 80 | | | | |
| 81 | | | | | 81 | | | | |
| 82 | | | | | 82 | | | | |
| 83 | 1.013 | 34.4 | + | 5 | 83 | 1.025 | 57.64 | + | 5 |
| 84 | | | | | 84 | | | | |
| 85 | | | | | 85 | | | | |
| 86 | | | | | 86 | | | | |
| 87 | | | | | 87 | | | | |
| 88 | | | | | 88 | | | | |
| 89 | | | | | 89 | | | | |
| 90 | 1.011 | 53.56 | + | 5 | 90 | 1.038 | 66.64 | + | 5 |
| 91 | | | | | 91 | | | | |
| 92 | | | | | 92 | | | | |
| 93 | | | | | 93 | | | | |
| 94 | | | | | 94 | | | | |
| 95 | | | | | 95 | | | | |
| 96 | | | | | 96 | | | | |
| 97 | | 80.6 | | | 97 | 1.033 | 95.04 | + | 4 |
| 98 | 1.009 | | + | 5 | 98 | | | | |
| 99 | | | | | 99 | | | | |
| 100 | | | | | 100 | | | | |
| 101 | | | | | 101 | | | | |
| 102 | | | | | 102 | | | | |
| 103 | | | | | 103 | | | | |
| 104 | 1.015 | 229.92 | + | 5 | 104 | 1.024 | 154.24 | + | 3 |
| 105 | | | | | 105 | | | | |
| 106 | | | | | 106 | | | | |
| 107 | | | | | 107 | | | | |
| 108 | | | | | 108 | | | | |
| 109 | | | | | 109 | | | | |
| 110 | | | | | 110 | | | | |
| 111 | 1.012 | 283.2 | + | 3 | 111 | 1.026 | 202.32 | − | 2 |
| 112 | | | | | 112 | | | | |
| 113 | | | | | 113 | | | | |
| 114 | | | | | 114 | | | | |
| 115 | | | | | 115 | | | | |
| 116 | | | | | 116 | | | | |
| 117 | | | | | 117 | | | | |
| 118 | 1.017 | 344.48 | + | 3 | 118 | 1.033 | 241.36 | − | 1 |
| 119 | | | | | 119 | | | | |
| 120 | | | | | 120 | | | | |
| 121 | | | | | 121 | | | | |
| 122 | | | | | 122 | | | | |
| 123 | | | | | 123 | | | | |
| 124 | | | | | 124 | | | | |
| 125 | 1.009 | 416 | ± | 2 | 125 | | 302.72 | − | 1 |
| 126 | | | | | 126 | 1.023 | | − | 1 |
| 127 | | | | | 127 | | | | |
| 128 | | | | | 128 | | | | |
| 129 | | | | | 129 | | | | |
| 130 | | | | | 130 | | | | |
| 131 | | | | | 131 | | | | |
| 132 | 1.015 | 368.64 | ± | 2 (2-3) | 132 | 1.038 | 284.4 | − | 1 |
| 133 | | | | | 133 | | | | |
| 134 | | | | | 134 | | | | |
| 135 | | | | | 135 | | | | |
| 136 | | | | | 136 | | | | |
| 137 | | | | | 137 | | | | |
| 138 | | | | | 138 | | | | |
| 139 | 1.014 | 343.52 | − | 2 | 139 | 1.04 | 245.68 | − | 1 |
| 140 | | | | | 140 | | | | |
| 141 | | | | | 141 | | | | |
| 142 | | | | | 142 | | | | |
| 143 | | | | | 143 | | | | |
| 144 | | | | | 144 | | | | |
| 145 | | | | | 145 | | | | |
| 146 | 1.013 | 225.60 | − | 2 | 146 | 1.038 | 235.68 | − | 1 |
| 147 | | | | | 147 | | | | |
| 148 | | | | | 148 | | | | |
| 149 | | | | | 149 | | | | |
| 150 | | | | | 150 | | | | |
| 151 | | | | | 151 | | | | |
| 152 | | | | | 152 | | | | |
| 153 | | 86.64 | − | 1 | Birth | 1.021 | | − | 1 |
| 154 | 1.016 | | − | 1 | 154 | | | | |
| Birth | | | | | 155 | | | | |
| 156 | | | | | 156 | | | | |
| 157 | | | | | 157 | | | | |
| 158 | | | | | 158 | | | | |
| 159 | | | | | 159 | | | | |
| 160 | | | | | 160 | | | | |
| 161 | | | | | 161 | | | | |
| 162 | | | | | 162 | | | | |
| 163 | | | | | 163 | | | | |
| 164 | | | | | 164 | | | | |
| 165 | | | | | 165 | | | | |

| I6142F | | | | | I6342F | | | | |
|---|---|---|---|---|---|---|---|---|---|
| Pregnancy date | *USG | **Progesterone (ng/mL) | Dual Checker Decision | Dual Checker Score | Pregnancy date | *USG | **Progesterone (ng/mL) | Dual Checker Decision | Dual Checker Score |
| Birth | | | | | Birth | | | | |
| 1 | 1.033 | | − | 1 | 1 | 1.041 | | − | 1 |
| 2 | 1.037 | | − | 1 | 2 | | | − | 1 |
| 3 | 1.039 | | − | 2 | 3 | 1.041 | | − | 2 |
| 4 | 1.038 | | − | 1 | 4 | 1.037 | | − | 1 |
| 5 | | | − | 1 | 5 | 1.039 | | − | 1 |
| 6 | | | − | 1 | 6 | 1.021 | | − | 1 |
| 7 | | 1.58 | − | 1 | 7 | 1.033 | 0.94 | − | 1 |
| 8 | 1..028 | | − | 1 | 8 | 1.04 | 5.32 | − | 1 |
| 9 | 1.039 | | − | 1 | 9 | | | − | 1 |
| 10 | 1.033 | | − | 1 | 10 | 1.039 | | − | 1 |
| 11 | 1.037 | 2.22 | − | 1 | 11 | 1.041 | 0.3 | + | 4 |
| 12 | | | − | 1 | 12 | 1.038 | | + | 3 |
| 13 | 1.028 | | − | 2 | 13 | 1.044 | 11.96 | ± | 2 |
| 14 | 1.028 | | + | 3 | 14 | 1.039 | | ± | 2 |
| 15 | 1.039 | 13.14 | + | 3 | 15 | 1.035 | 29.32 | ± | 3 |
| 16 | 1.03 | | ± | 2 | 16 | | | − | 2 |
| 17 | 1.038 | | ± | 2 | 17 | | | − | 2 |
| 18 | 1.038 | | − | 2 | 18 | | | ± | 2 |
| 19 | | | − | 1 | 19 | 1.038 | | − | 1 |
| 20 | 1.034 | | − | 2 | 20 | 1.022 | | − | 2 |
| 21 | 1.037 | | − | 1 | 21 | 1.03 | | − | 1 |
| 22 | 1.03 | | − | 1 | 22 | 1.044 | | − | 2 |
| 23 | 1.032 | | − | 2 | 23 | | | − | 2 |
| 24 | 1.029 | 105.72 | − | 2 | 24 | | | − | 1 |
| 25 | | | − | 1 | 25 | 1.035 | | − | 2 |
| 26 | | | − | 1 | 26 | 1.039 | 43.56 | + | 3 |
| 27 | | | − | 1 | 27 | 1.03 | | + | 3 |
| 28 | | | − | 1 | 28 | 1.033 | | + | 4 |
| 29 | 1.031 | | − | 2 | 29 | | | + | 3 |
| 30 | 1.031 | | − | 2 | 30 | | | + | 4 |
| 31 | 1.03 | 91.72 | + | 3 | 31 | | | + | 4 |
| 32 | 1.035 | | + | 3 | 32 | 1.03 | | + | 4 |
| 33 | | | + | 3 | 33 | 1.032 | 47.36 | + | 4 |
| 34 | 1.032 | | + | 4 | 34 | | | | |
| 35 | 1.034 | | + | 4 | 35 | | | | |
| 36 | × | | + | 4 | 36 | | | | |
| 37 | 1.031 | | + | 4 | 37 | | | | |
| 38 | 1.031 | 88.04 | + | 4 | 38 | | | | |
| 39 | | | | | 39 | | | | |
| 40 | | | | | 40 | 1.039 | 48.36 | + | 5 |
| 41 | | | | | 41 | | | | |
| 42 | | | | | 42 | | | | |
| 43 | | | | | 43 | | | | |
| 44 | | | | | 44 | | | | |
| 45 | 1.034 | 86.72 | + | 4 | 45 | | | | |
| 46 | | | | | 46 | | | | |
| 47 | | | | | 47 | 1.034 | 49.84 | + | 5 |
| 48 | | | | | 48 | | | | |
| 49 | | | | | 49 | | | | |
| 50 | | | | | 50 | | | | |
| 51 | | | | | 51 | | | | |
| 52 | 1.034 | 77.48 | + | 4 | 52 | | | | |
| 53 | | | | | 53 | | | | |
| 54 | | | | | 54 | 1.036 | 34.48 | + | 5 |
| 55 | | | | | 55 | | | | |
| 56 | | | | | 56 | | | | |
| 57 | | | | | 57 | | | | |
| 58 | | | | | 58 | | | | |
| 59 | 1.027 | 122.76 | + | 5 | 59 | | | | |
| 60 | | | | | 60 | | | | |
| 61 | | | | | 61 | 1.037 | 41.04 | + | 5 |
| 62 | | | | | 62 | | | | |
| 63 | | | | | 63 | | | | |
| 64 | | | | | 64 | | | | |
| 65 | | | | | 65 | | | | |
| 66 | 1.023 | 110.88 | + | 5 | 66 | | | | |
| 67 | | | | | 67 | | | | |
| 68 | | | | | 68 | 1.023 | 38.04 | + | 5 |
| 69 | | | | | 69 | | | | |
| 70 | | | | | 70 | | | | |
| 71 | | | | | 71 | | | | |
| 72 | | | | | 72 | | | | |
| 73 | 1.022 | 106.52 | + | 5 | 73 | | | | |
| 74 | | | | | 74 | | | | |
| 75 | | | | | 75 | 1.027 | 42.88 | + | 5 |
| 76 | | | | | 76 | | | | |
| 77 | | | | | 77 | | | | |
| 78 | | | | | 78 | | | | |
| 79 | | | | | 79 | | | | |
| 80 | 1.026 | 108.64 | + | 5 | 80 | | | | |
| 81 | | | | | 81 | | | | |
| 82 | | | | | 82 | 1.028 | 46.28 | + | 5 |
| 83 | | | | | 83 | | | | |
| 84 | | | | | 84 | | | | |
| 85 | | | | | 85 | | | | |
| 86 | | | | | 86 | | | | |
| 87 | 1.025 | 109.2 | + | 5 | 87 | | | | |
| 88 | | | | | 88 | | | | |
| 89 | | | | | 89 | 1.028 | 46.2 | + | 5 |
| 90 | | | | | 90 | | | | |
| 91 | | | | | 91 | | | | |
| 92 | | | | | 92 | | | | |
| 93 | | | | | 93 | | | | |
| 94 | 1.026 | 98.44 | + | 5 | 94 | | | | |
| 95 | | | | | 95 | | | | |
| 96 | | | | | 96 | 1.021 | 56.16 | + | 5 |
| 97 | | | | | 97 | | | | |
| 98 | | | | | 98 | | | | |
| 99 | | | | | 99 | | | | |
| 100 | | | | | 100 | | | | |
| 101 | 1.008 | 123.48 | + | 4 | 101 | | | | |
| 102 | | | | | 102 | | | | |
| 103 | | | | | 103 | 1.023 | 100.6 | + | 4 |
| 104 | | | | | 104 | | | | |
| 105 | | | | | 105 | | | | |
| 106 | | | | | 106 | | | | |
| 107 | | | | | 107 | | | | |
| 108 | 1.009 | 177.84 | + | 3 | 108 | | | | |
| 109 | | | | | 109 | | | | |
| 110 | | | | | 110 | 1.031 | 149.72 | + | 3 |
| 111 | | | | | 111 | | | | |
| 112 | | | | | 112 | | | | |
| 113 | | | | | 113 | | | | |
| 114 | | | | | 114 | | | | |
| 115 | 1.016 | 218.88 | − | 2 | 115 | | | | |
| 116 | | | | | 116 | | | | |
| 117 | | | | | 117 | 1.033 | 159.76 | − | 2 |
| 118 | | | | | 118 | | | | |
| 119 | | | | | 119 | | | | |
| 120 | | | | | 120 | | | | |
| 121 | | | | | 121 | | | | |
| 122 | 1.031 | 258 | − | 1 | 122 | | | | |
| 123 | | | | | 123 | | | | |
| 124 | | | | | 124 | 1.031 | 225.44 | − | 1 |
| 125 | | | | | 125 | | | | |
| 126 | | | | | 126 | | | | |
| 127 | | | | | 127 | | | | |
| 128 | | | | | 128 | | | | |
| 129 | 1.025 | 245.92 | − | 1 | 129 | | | | |
| 130 | | | | | 130 | | | | |
| 131 | | | | | 131 | 1.029 | 266.72 | − | 1 |
| 132 | | | | | 132 | | | | |
| 133 | | | | | 133 | | | | |
| 134 | | | | | 134 | | | | |
| 135 | | | | | 135 | | | | |
| 136 | 1.037 | 200.8 | − | 1 | 136 | | | | |
| 137 | | | | | 137 | | | | |
| 138 | | | | | 138 | 1.032 | 289.68 | − | 1 |
| 139 | | | | | 139 | | | | |
| 140 | | | | | 140 | | | | |
| 141 | | | | | 141 | | | | |
| 142 | | | | | 142 | | | | |
| 143 | 1.035 | 301.12 | − | 1 | 143 | | | | |
| 144 | | | | | 144 | | | | |
| 145 | | | | | 145 | 1.035 | 197.76 | − | 1 |
| 146 | | | | | 146 | | | | |
| 147 | | | | | 147 | | | | |
| 148 | | | | | 148 | | | | |
| 149 | | | | | 149 | | | | |
| 150 | 1.026 | 98.56 | − | 1 | 150 | | | | |
| 151 | | | | | 151 | | | | |
| 152 | | | | | 152 | 1.04 | 178.72 | − | 1 |
| 153 | | | | | 153 | | | | |
| 154 | | | | | 154 | | | | |
| 155 | | | | | 155 | | | | |
| 156 | | | | | 156 | | | | |
| 157 | 1.028 | 57.68 | − | 2 | Birth | | | | |
| 158 | | | | | 158 | | | | |
| Birth | | | | | 159 | 1.043 | 7.1 | − | 2 |
| 160 | | | | | 160 | | | | |
| 161 | | | | | 161 | | | | |
| 162 | | | | | 162 | | | | |
| 163 | | | | | 163 | | | | |
| 164 | 1.017 | 1.9 | − | 1 | 164 | | | | |
| 165 | | | | | 165 | | | | |

| I6193F | | | | | I5592F | | | | |
|---|---|---|---|---|---|---|---|---|---|
| Pregnancy date | *USG | **Progesterone (ng/mL) | Dual Checker Decision | Dual Checker Score | Pregnancy date | *USG | **Progesterone (ng/mL) | Dual Checker Decision | Dual Checker Score |
| Birth | | | | | Birth | | | | |
| 1 | 1.039 | | − | 1 | 1 | 1.034 | | − | 1 |
| 2 | 1.043 | | − | 1 | 2 | 1.034 | | − | 1 |
| 3 | 1.017 | | − | 1 | 3 | 1.027 | | − | 1 |
| 4 | 1.039 | | − | 1 | 4 | 1.025 | | − | 1 |
| 5 | 1.043 | 2.04 | − | 1 | 5 | 1.031 | | − | 1 |
| 6 | | | − | 1 | 6 | 1.028 | | − | 1 |
| 7 | | | − | 1 | 7 | 1.031 | 6.32 | − | 1 |
| 8 | | | − | 1 | 8 | 1.032 | | − | 1 |
| 9 | 1.037 | 3.78 | − | 1 | 9 | 1.032 | | − | 1 |
| 10 | 1.038 | | + | 3 | 10 | 1.026 | 19.58 | + | 3 |
| 11 | 1.033 | | − | 2 | 11 | 1.024 | | + | 3 |
| 12 | 1.034 | 21.26 | − | 1 | 12 | 1.031 | 27.7 | − | 1 |
| 13 | | | − | 2 | 13 | 1.027 | | − | 2 |
| 14 | | | − | 1 | 14 | 1.033 | | − | 1 |
| 15 | 1.038 | | − | 2 | 15 | 1.035 | | − | 2 |
| 16 | 1.038 | 42.52 | − | 2 | 16 | 1.032 | | − | 2 |
| 17 | 1.028 | | − | 1 | 17 | 1.028 | | − | 2 |
| 18 | 1.033 | | − | 1 | 18 | 1.032 | 100.28 | − | 1 |
| 19 | | | − | 1 | 19 | 1.026 | | − | 1 |
| 20 | | | − | 1 | 20 | 1.034 | | − | 1 |
| 21 | | | − | 1 | 21 | 1.031 | | − | 1 |
| 22 | 1.037 | | − | 1 | 22 | 1.034 | | − | 1 |
| 23 | 1.028 | 24.48 | − | 1 | 23 | 1.032 | | − | 1 |
| 24 | 1.03 | | − | 1 | 24 | 1.033 | | − | 1 |
| 25 | 1.032 | | − | 1 | 25 | 1.03 | 143.2 | − | 2 |
| 26 | 1.034 | | ± | 2 | 26 | 1.03 | | − | 2 |
| 27 | 1.031 | | − | 2 | 27 | 1.029 | | ± | 3 |
| 28 | 1.028 | | ± | 2 | 28 | 1.028 | | ± | 2 |
| 29 | 1.027 | | + | 3 | 29 | 1.032 | | ± | 3 |
| 30 | 1.03 | 85.76 | + | 3 | 30 | 1.037 | | + | 3 |
| 31 | 1.015 | | + | 3 | 31 | 1.032 | | + | 4 |
| 32 | 1.031 | | + | 3 | 32 | 1.035 | 124.2 | + | 3 |
| 33 | | | | | 33 | | | | |
| 34 | | | | | 34 | | | | |
| 35 | | | | | 35 | | | | |
| 36 | | | | | 36 | | | | |
| 37 | 1.03 | 76.52 | + | 3 | 37 | | | | |
| 38 | | | | | 38 | | | | |
| 39 | | | | | 39 | 1.035 | 102.16 | + | 4 |
| 40 | | | | | 40 | | | | |
| 41 | | | | | 41 | | | | |
| 42 | | | | | 42 | | | | |
| 43 | | | | | 43 | | | | |
| 44 | 1.028 | 80.84 | + | 4 | 44 | | | | |
| 45 | | | | | 45 | | | | |
| 46 | | | | | 46 | 1.024 | 108.56 | + | 4 |
| 47 | | | | | 47 | | | | |
| 48 | | | | | 48 | | | | |
| 49 | | | | | 49 | | | | |
| 50 | | | | | 50 | | | | |
| 51 | 1.029 | 106.56 | + | 4 | 51 | | | | |
| 52 | | | | | 52 | | | | |
| 53 | | | | | 53 | 1.034 | 105.8 | + | 4 |
| 54 | | | | | 54 | | | | |
| 55 | | | | | 55 | | | | |
| 56 | | | | | 56 | | | | |
| 57 | | | | | 57 | | | | |
| 58 | 1.031 | 97.52 | + | 5 | 58 | | | | |
| 59 | | | | | 59 | | | | |
| 60 | | | | | 60 | 1.031 | 127.08 | + | 5 |
| 61 | | | | | 61 | | | | |
| 62 | | | | | 62 | | | | |
| 63 | | | | | 63 | | | | |
| 64 | | | | | 64 | | | | |
| 65 | 1.022 | 120.4 | + | 5 | 65 | | | | |
| 66 | | | | | 66 | | | | |
| 67 | | | | | 67 | 1.033 | 142.68 | + | 5 |
| 68 | | | | | 68 | | | | |
| 69 | | | | | 69 | | | | |
| 70 | | | | | 70 | | | | |
| 71 | | | | | 71 | | | | |
| 72 | 1.017 | 114.76 | + | 5 | 72 | | | | |
| 73 | | | | | 73 | | | | |
| 74 | | | | | 74 | 1.026 | 118.28 | + | 5 |
| 75 | | | | | 75 | | | | |
| 76 | | | | | 76 | | | | |
| 77 | | | | | 77 | | | | |
| 78 | | | | | 78 | | | | |
| 79 | 1.021 | 115.4 | + | 5 | 79 | | | | |
| 80 | | | | | 80 | | | | |
| 81 | | | | | 81 | 1.027 | 100.56 | + | 5 |
| 82 | | | | | 82 | | | | |
| 83 | | | | | 83 | | | | |
| 84 | | | | | 84 | | | | |
| 85 | | | | | 85 | | | | |
| 86 | 1.035 | 110.96 | + | 5 | 86 | | | | |
| 87 | | | | | 87 | | | | |
| 88 | | | | | 88 | 1.032 | 77.76 | + | 5 |
| 89 | | | | | 89 | | | | |
| 90 | | | | | 90 | | | | |
| 91 | | | | | 91 | | | | |
| 92 | | | | | 92 | | | | |
| 93 | 1.034 | 123.2 | + | 4 | 93 | | | | |
| 94 | | | | | 94 | | | | |
| 95 | | | | | 95 | 1.021 | 134.24 | + | 5 |
| 96 | | | | | 96 | | | | |
| 97 | | | | | 97 | | | | |
| 98 | | | | | 98 | | | | |
| 99 | | | | | 99 | | | | |
| 100 | 1.027 | 157.48 | + | 4 | 100 | | | | |
| 101 | | | | | 101 | | | | |
| 102 | | | | | 102 | 1.013 | 118.48 | + | 4 |
| 103 | | | | | 103 | | | | |
| 104 | | | | | 104 | | | | |
| 105 | | | | | 105 | | | | |
| 106 | | | | | 106 | | | | |
| 107 | 1.043 | 116.4 | − | 2 | 107 | | | | |
| 108 | | | | | 108 | | | | |
| 109 | | | | | 109 | 1.022 | 127.92 | + | 3 |
| 110 | | | | | 110 | | | | |
| 111 | | | | | 111 | | | | |
| 112 | | | | | 112 | | | | |
| 113 | | | | | 113 | | | | |
| 114 | 1.027 | 132.56 | − | 2 | 114 | | | | |
| 115 | | | | | 115 | | | | |
| 116 | | | | | 116 | 1.018 | 156.72 | ± | 2 |
| 117 | | | | | 117 | | | | |
| 118 | | | | | 118 | | | | |
| 119 | | | | | 119 | | | | |
| 120 | | | | | 120 | | | | |
| 121 | 1.01 | 165.2 | − | 1 | 121 | | | | |
| 122 | | | | | 122 | | | | |
| 123 | | | | | 123 | 1.018 | 182.96 | − | 1 |
| 124 | | | | | 124 | | | | |
| 125 | | | | | 125 | | | | |
| 126 | | | | | 126 | | | | |
| 127 | | | | | 127 | | | | |
| 128 | 1.02 | 190.32 | − | 1 | 128 | | | | |
| 129 | | | | | 129 | | | | |
| 130 | | | | | 130 | 1.019 | 188.64 | − | 1 |
| 131 | | | | | 131 | | | | |
| 132 | | | | | 132 | | | | |
| 133 | | | | | 133 | | | | |
| 134 | | | | | 134 | | | | |
| 135 | 1.012 | 181.92 | − | 1 | 135 | | | | |
| 136 | | | | | 136 | | | | |
| 137 | | | | | 137 | 1.019 | 182.8 | − | 1 |
| 138 | | | | | 138 | | | | |
| 139 | | | | | 139 | | | | |
| 140 | | | | | 140 | | | | |
| 141 | | | | | 141 | | | | |
| 142 | 1.009 | 173.36 | − | 1 | 142 | | | | |
| 143 | | | | | 143 | | | | |
| 144 | | | | | 144 | 1.023 | 235.2 | − | 1 |
| 145 | | | | | 145 | | | | |
| 146 | | | | | 146 | | | | |
| 147 | | | | | 147 | | | | |
| 148 | | | | | 148 | | | | |
| 149 | 1.039 | 111.6 | − | 1 | 149 | | | | |
| 150 | | | | | 150 | | | | |
| 151 | | | | | 151 | 1.01 | 380 | − | 1 |
| 152 | | | | | 152 | | | | |
| 153 | | | | | 153 | | | | |
| Birth | | | | | 154 | | | | |
| 155 | | | | | 155 | | | | |
| 156 | 1.036 | 1.48 | − | 1 | 156 | | | | |
| 157 | | | | | Birth | | | | |
| 158 | | | | | 158 | 1.031 | 1.58 | − | 1 |
| 159 | | | | | 159 | | | | |
| 160 | | | | | 160 | | | | |
| 161 | | | | | 161 | | | | |
| 162 | | | | | 162 | | | | |
| 163 | | | | | 163 | | | | |
| 164 | | | | | 164 | | | | |
| 165 | | | | | 165 | | | | |

*USG: urine specific gravity

**Progesterone concentration was mesured by using plasma

**Supplementary table S5: The result for CG assay using cynomolgus monkey**

| Cynomolgus monkey ID | ultrasonographic examination | Period for mating | Estradiol (pg/mL) | Progesterone (ng/mL) | *Timing for CG assay | CG test kit |
|---|---|---|---|---|---|---|
| 1411407053 | ＋ | AI | 42.1 | 4.67 | day30 | － |
| 1411906047 | － | for 3 days | 200.5 | ＜L | day23 | － |
| 1411910111 | － | for 3 days | 25.7 | 4.12 | day20 | － |
| 1412005031 | ＋ | for 7 days | 83.9 | 6.2 | day24 | － |
| 1511805015 | － | for 3 days | 58.9 | 0.16 | day21 | － |
| 1512006025 | ＋ | for 3 days | 56.7 | 14.25 | day23 | － |
| 1512006030 | ＋ | for 3 days | 99.2 | 9.34 | day24 | － |
| 1512008057 | ＋ | for 3 days | 61.8 | 61.8 | day25 | － |
| 1512011077 | － | for 3 days | ＜L | 0.15 | day22 | － |
| 1611810004 | － | for 3 days | 78.3 | 0.24 | day21 | － |

*: This timing is the days from mating.

**Supplementary table S6: The obtained marmoset CG mAbs**

#### Anti-CGα

| No. | Clone | Subclass |
|---|---|---|
| 1 | 1F1 | — |
| 2 | 5H3 | IgG 1 κ |
| 3 | 46A4 | IgG 2a κ |
| 4 | 46F7 | IgG 1 κ |
| 5 | 52B7 | IgG 1 κ |
| 6 | 53F3 | IgG 1 κ |
| 7 | 54B9 | IgG 1 κ |
| 8 | 56C7 | IgG 1 κ |
| 9 | 56G2 | IgG 1 κ |
| 10 | 57C7 | IgG 1 κ |
| 11 | 58C8 | IgG 1 κ |
| 12 | 58F10 | IgG 1 κ |
| 13 | 59E1 | IgG 2a κ |
| 14 | 65C10 | IgG 2a κ |

#### Anti-CGβ

| No. | Clone | Subclass |
|---|---|---|
| 15 | 4E3 | IgG 2a κ |
| 16 | 8B1 | IgG 1 κ |
| 17 | 9E11 | IgG 1 κ |
| 18 | 13B6 | IgG 1 κ |
| 19 | 13D7 | IgG 2a λ |
| 20 | 29C2 | IgG 1 κ |
| 21 | 31D9 | IgG 2a κ |
| 22 | 33G9 | IgG 1 κ |

#### Supplementary table S7: The result of sandwich ELISA for the development of CG immunochromatography

| | | Anti-CGβ | | | | | | | | | | | | | | | |
|---|---|---|---|---|---|---|---|---|---|---|---|---|---|---|---|---|---|
| | | 4E3 | | 8B1 | | 9E11 | | 13B6 | | 13D7 | | 29C2 | | 31D9 | | 33G9 | |
| Concentration of CG (AU/mL) | | 0 | 10 | 0 | 10 | 0 | 10 | 0 | 10 | 0 | 10 | 0 | 10 | 0 | 10 | 0 | 10 |
| Anti-CGα | 5H3 | -0.002 | 0.097 | -0.014 | 0.197 | -0.002 | 0.380 | -0.005 | 0.836 | 0 | 0.426 | -0.021 | 0.598 | 0.007 | 0.364 | 0.003 | 1.019 |
| | 59E1 | 0.003 | 0.139 | 0.045 | 0.306 | 0.078 | 0.367 | 0.025 | 0.527 | 0.021 | 0.360 | 0.003 | 0.786 | 0.049 | 0.367 | 0.084 | 0.732 |
| | 46A4 | 0.042 | 0.206 | 0.057 | 0.320 | 0.065 | 0.540 | 0.023 | 1.057 | 0.035 | 0.589 | -0.018 | 0.820 | 0.054 | 0.568 | 0.053 | 1.224 |
| | 65C10 | 0.043 | 0.204 | 0.017 | 0.310 | 0.027 | 0.528 | 0.003 | 1.055 | 0.023 | 0.947 | -0.029 | 0.947 | 0.066 | 0.630 | 0.074 | 1.319 |
| | 52B7 | 0.007 | 0.190 | 0.000 | 0.384 | 0.056 | 0.731 | 0.014 | 1.485* | -0.022 | 0.780 | -0.022 | 1.132 | 0.033 | 0.747 | 0.035 | 1.719 |

*The red letter indicated the result of the best combination.
The low background and the high absorbance of 10 AU/mL were selected.
